# Ring Finger Protein 213 Assembles into a Sensor for ISGylated Proteins with Antimicrobial Activity

**DOI:** 10.1101/2021.06.03.446796

**Authors:** Fabien Thery, Lia Martina, Caroline Asselman, Heidi Repo, Yifeng Zhang, Koen Sedeyn, George D. Moschonas, Clara Bredow, Qi Wen Teo, Jingshu Zhang, Madeleine Vessely, Kevin Leandro, Denzel Eggermont, Delphine De Sutter, Katie Boucher, Tino Hochepied, Nele Festjens, Nico Callewaert, Xavier Saelens, Bart Dermaut, Klaus-Peter Knobeloch, Antje Beling, Sumana Sanyal, Lilliana Radoshevich, Sven Eyckerman, Francis Impens

## Abstract

ISG15 is an interferon-stimulated, ubiquitin-like protein that can conjugate to substrate proteins (ISGylation) to counteract microbial infection, but the underlying mechanisms remain elusive. Here, we used a viral-like particle trapping technology to identify ISG15-binding proteins and discovered Ring Finger Protein 213 (RNF213) as an ISG15 interactor and cellular sensor of ISGylated proteins. RNF213 is a poorly-characterized, interferon-induced megaprotein that is frequently mutated in Moyamoya disease, a rare cerebrovascular disorder. We found that interferon induces ISGylation and oligomerization of RNF213 on lipid droplets, where it acts as a sensor for ISGylated proteins. We showed that RNF213 has broad antimicrobial activity in vitro and in vivo, counteracting infection with Listeria monocytogenes, herpes simplex virus 1 (HSV-1), human respiratory syncytial virus (RSV) and coxsackievirus B3 (CVB3), and we observed a striking co-localization of RNF213 with intracellular bacteria. Together, our findings provide novel molecular insights into the ISGylation pathway and reveal RNF213 as a key antimicrobial effector.

## INTRODUCTION

ISG15 is a ubiquitin-like (UBL) protein with antimicrobial activity. Similar to ubiquitin, ISG15 conjugates via its C-terminus to lysine residues of substrate proteins in a process called ISGylation, which is mediated by an E1 enzyme, UBE1L, an E2 enzyme, UBCH8, and three known E3 enzymes, HHARI, TRIM25 and HERC5 ^1–5^. ISG15 and its conjugation machinery are strongly upregulated by Type I and III interferon, viral nucleic acids ^1^, bacterial DNA ^6^ and lipopolysaccharide (LPS) ^7^. ISG15 has potent antiviral effects both *in vitro* and *in vivo* ^8^. In fact, mice which lack ISG15 are unable to control various viral pathogens including clinically relevant etiologic agents such as Influenza ^9^, human respiratory syncytial virus ^10^ and coxsackievirus ^11,12^. ISG15’s crucial antiviral role is bolstered by the variety of viral evasion strategies targeting the ISGylation pathway, either by interfering with ISG15 conjugation ^1,13^ or by expressing ISG15- specific proteases that lead to reversible ^14–17^ or irreversible deconjugation ^18^. In addition to its antiviral role, ISG15 can also act as an antibacterial effector against intracellular bacterial pathogens such as *Listeria monocytogenes* ^6^ and *Mycobacterium tuberculosis* ^19^ and it can restrict the intracellular eukaryotic pathogen *Toxoplasma gondii* ^20^. Despite its broad antimicrobial role, the molecular mechanisms by which ISG15 modification is sensed and how it protects against microbial infections remain largely undefined. One model posits that the antiviral function of ISG15 is based on the localization of the E3 ligase HERC5 at the ribosome. According to this model, proteins are co-translationally modified by ISG15 during infection, thereby interfering with the function of newly translated viral proteins in a non-specific manner ^21^. However, the model does not predict what subsequently happens to ISGylated proteins in the cell and how they are recognized and trafficked.

Unlike ubiquitin, ISG15 does not solely exert its antimicrobial function by covalently conjugating to target proteins. It can also be secreted and act as a cytokine ^22–24^ and can non-covalently interact with viral and host proteins ^25^. Relatively little is known about what interaction partners bind ISG15 and how the functions of these interactions contribute to host responses to pathogens. Indeed, only a few host proteins have been reported to non-covalently bind to ISG15, mainly in its free form. One of the best-characterized ISG15- binding proteins is USP18, the predominant ISG15 deconjugating protease in human and mice ^26,27^. In addition to its enzymatic activity, USP18 functions as a major negative regulator of type 1 interferon (IFN-I) signaling by binding to one of the subunits of the interferon α/β receptor (IFNAR2) ^28–30^. In order to prevent IFN-I over-amplification in humans, USP18 needs to be stabilized by binding to free ISG15 in a non-covalent manner. Hence, patients who lack ISG15 expression due to a frame-shift mutation have a strong upregulation of the IFN-I pathway leading to an Aicardi-Goutières-like interferonopathy ^31^. In contrast to *isg15*-deficient mice, these patients do not display enhanced susceptibility to viral infection, instead they are susceptible to bacterial infection including clearance of the attenuated tuberculosis vaccine, Bacille Calmette-Guerin (BCG) ^32^. In addition to USP18, a few other proteins have been reported to interact with ISG15 in a non-covalent manner. Free intracellular ISG15 can bind to leucine-rich repeat-containing protein 25 (LRRC25) and mediate the autophagic degradation of retinoic acid-inducible gene I protein (RIG-I/DDX58) ^33^. Upon forced overexpression, p62 and HDAC6 were further shown to interact with the C-terminal LRLRGG sequence of ISG15 (and ISGylated proteins) ^34^. Free ISG15 also interacts with the E3 ligase NEDD4 and ISG15 overexpression disrupts NEDD4 ubiquitination activity thus blocking the budding of Ebola virus-like particles ^35,36^. Likewise, free ISG15 was reported to interact in a non-covalent manner with the Hypoxia inducible factor 1α (HIF1α), preventing its dimerization and downstream signaling ^37^.

ISG15 has not been studied as extensively as ubiquitin or SUMO for which interacting domains or motifs have been reported, thus to date no such domains or motifs have been described for ISG15. Although affinity purification mass spectrometry (AP-MS) and yeast-two-hybrid (Y2H) screens were performed with ISG15 as bait, these interactome screens were hampered by the inherently limited binding surface of such a small protein and the stringency of yeast two-hybrid, which would preclude low affinity interactions ^38–41^. Therefore, we endeavored to identify non-covalent interaction partners of ISG15 using a recently-developed approach named Virotrap, which is based on capturing protein complexes within virus-like particles (VLPs) that bud from mammalian cells ^42^. Virotrap is a cell lysis-free method and thus preserves existing protein complexes, and, importantly, this method is uniquely suited to capture weak and transient cytosolic interactions that otherwise do not survive protein complex purification. Using this technology, we systematically mapped the non-covalent interactome of ISG15 in human cells. We identified a rich interactome but for further study focused on a very-large protein, called Ring Finger Protein 213 (RNF213), mutations in which predisposes patients to early cerebrovascular events. Further biochemical evaluation confirmed the selective binding of RNF213 to ISG15 and showed that IFN-I signaling induces RNF213 oligomerization into a sensor for ISGylated proteins associated with lipid droplets. Finally, *in vitro* and *in vivo* loss and gain of function experiments showed that RNF213 is a pivotal, broadly acting microbial restriction factor that colocalizes to the surface of intracellular bacteria.

## RESULTS

### Identification of RNF213 as an ISG15-binding protein by Virotrap

To investigate the non-covalent interactome of ISG15 and cytosolic sensors for ISG15 modification, we used Virotrap, a recently developed MS-based approach to map protein-protein interactions ^42^. Briefly, the sequence of mature ISG15 (ending on -LRLRGG) was genetically fused via its N-terminus to the GAG protein of HIV-1 and expressed in HEK293T cells treated with IFN-I. Expression of the GAG-ISG15 fusion protein led to self-assembly of VLPs internally coated with ISG15. After budding from the cells and capturing ISG15 interaction partners, VLPs were collected from the cell culture supernatants and used for MS-based protein identification (Fig. 1A). To distinguish specific ISG15-binding proteins from non-specific (GAG-binding) proteins, the experiment was performed in a quantitative fashion comparing ISG15 VLPs with VLPs containing dihydrofolate reductase from *Escherichia coli* (eDHFR), a non-specific bait protein with no obvious interaction partners in human cells. In addition to ISG15, we identified RNF213 as the most enriched protein in the ISG15 VLPs, associated with high spectral and peptide counts (Fig. 1B, Table S1). Of note, among several less enriched proteins we also identified USP18, which is an ISG15 specific isopeptidase and thus a well-known interaction partner of ISG15 ^26,27^. To rule out any type of covalent interaction between GAG-ISG15 and RNF213, we repeated the Virotrap experiment using the non-conjugatable ISG15AA (with the c-terminal glycines mutated to alanines: LRLRAA) and precursor variants of ISG15, both with and without IFN-I treatment (Supplementary Fig. 1, Table S1). Without exception, in all conditions we identified RNF213 as the most enriched protein in the ISG15 VLPs. We also confirmed the presence of RNF213 in the ISG15 VLPs by western blotting (Fig. 1C).

**Figure 1.**
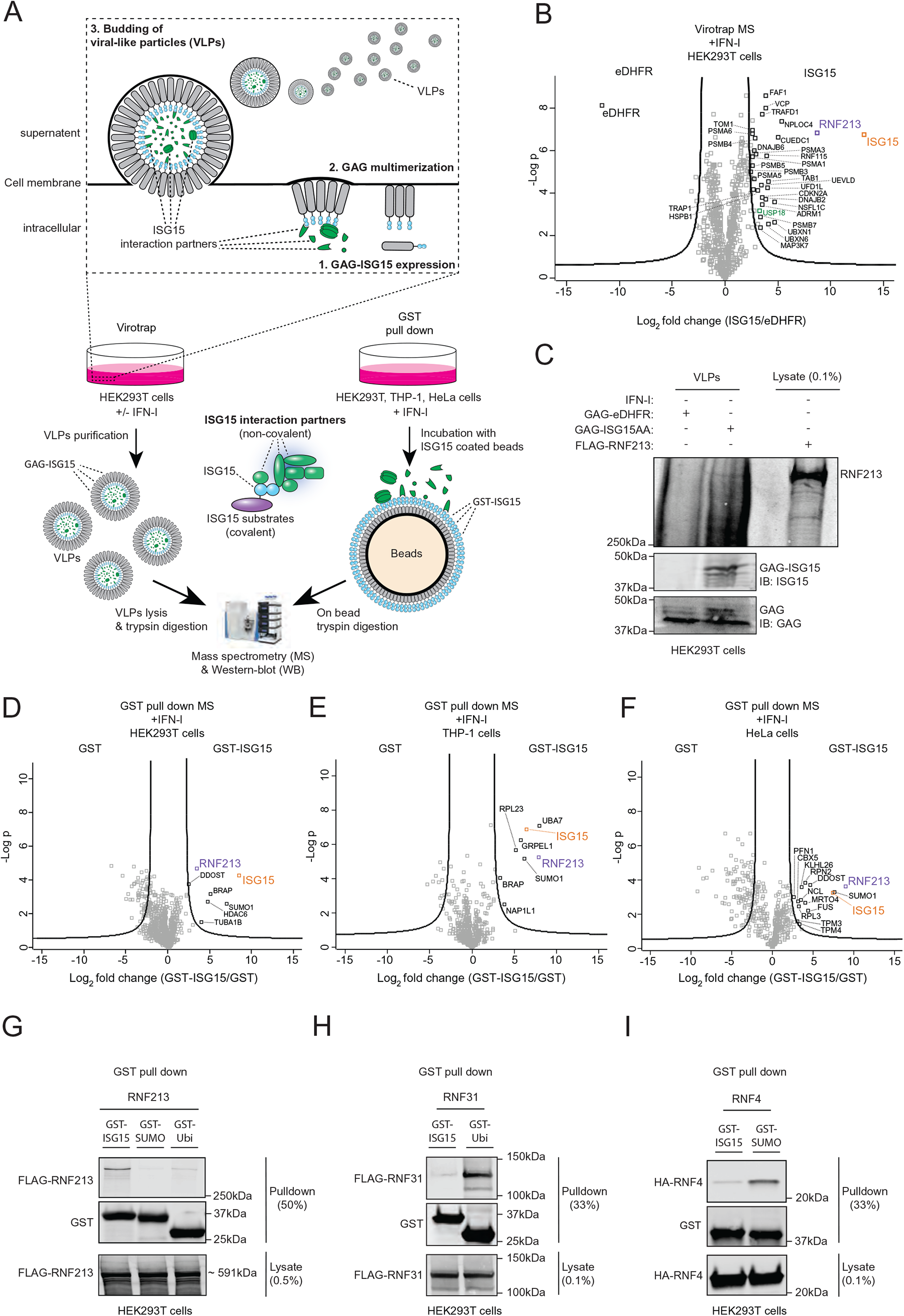
Identification of RNF213 as ISG15-binding protein. **(A)** Workflow showing the strategy for mapping ISG15 interaction partners. Virotrap and GST pulldown were employed as two orthogonal methods. In Virotrap, GAG-ISG15 was transiently expressed in HEK293T cells. 24 h post-transfection, cells were either treated with 10 ng/mL interferon-α for 24 h of left untreated. Budding virus-like particles (VLPs) containing ISG15 and its interaction partners were purified, lysed and digested into peptides prior LC-MS/MS analysis. In the GST pull down, glutathione beads were decorated with GST-ISG15 and mixed with a cellular lysate from HEK293T, HeLa, or THP-1 cells. Prior to lysis cells were treated for 24 h with 10 ng/mL interferon-α (HEK293T) or interferon-β (HeLa and THP- 1). Following on-bead digestion, the resulting peptides were analyzed by LC-MS/MS. **(B)** Volcano plot showing the result of a t-test to compare VLPs containing mature ISG15 versus VLPs containing dihydrofolate reductase from *Escherichia coli* (eDHFR) as negative control (n=4 replicates). The fold change (in log2) of each protein is shown on the x-axis, while the statistical significance (−log P-value) is shown on the y-axis. Proteins outside the curved lines represent specific ISG15 interaction partners. Proteins identified as ISG15 interaction partners in all virotrap screens, are annotated (n=29) and listed in Table S1**. (C)** VLPs containing ISG15 or eDHFR were collected and analyzed by immunoblot against RNF213, ISG15 and GAG. FLAG-RNF213 purified from a lysate of HEK293T cells was loaded as a positive control and confirmed the presence of an RNF213 band in the ISG15 VLPs. **(D-F)** Volcano plots comparing GST pull downs using GST-ISG15-coated beads with GST-coated beads as control in lysates of HEK293T, THP-1 and HeLa cells (n=3). Proteins significantly enriched in pull downs with ISG15- coated beads are annotated. **(G)** GST pull downs with ISG15, ubiquitin and SUMO followed by immunoblotting show binding of RNF213 to ISG15, but not to ubiquitin or SUMO. Beads coated with each UBL were mixed with a lysate of HEK293T cells expressing FLAG-RNF213 and bound proteins were analyzed by immunoblot against FLAG and GST. (**H-I**) Validation of GST pull down assays with ubiquitin and SUMO using RNF31 and RNF4 as known binders of these UBLs, respectively. Assays were performed as in (**G**).

In addition to RNF213, 28 other proteins were enriched in ISG15 VLPs under all conditions (Supplementary Fig. 1, Table S1). Following RNF213, the most notable among these identifications was the valosin-containing protein (VCP/p97). VCP is a hexameric AAA+ ATPase with a central role in the endoplasmic-reticulum-associated protein degradation (ERAD) pathway, where it associates with cofactor proteins to facilitate the proteasomal degradation of misfolded proteins ^43^. In addition, several cofactors of VCP were found to interact with ISG15 such as the VCP-UFD1-NPL4 protein complex as well as PLAA, UBXN6 and YOD1 ^44^, FAF1, NSFL1C, UBXN1 and RNF115 ^45–48^. We also detected several components of the proteasome complex enriched in ISG15 VLPs, including ADRM1 and several PSMA and PSMB subunits of the proteasome. Moreover, we identified TAB1 and TAK1 (MAP3K7) which are involved in innate immune signaling pathways as well as TRAFD1 ^49,50^.

We then performed an additional series of Virotrap-MS experiments in which only the N- or C-terminal domain of ISG15 was fused to GAG, alongside with full length ISG15 and eDHFR as positive and negative controls, respectively. None of the VLPs with the N- or C-terminal ubiquitin-like domain of ISG15 could enrich RNF213, suggesting that both domains are required for the interaction with RNF213 (Supplementary Fig. 2A-C). Like for RNF213, the majority of the 28 aforementioned ISG15 interaction partners also required both domains to interact (Supplementary Fig. 2D), suggesting that these proteins might interact indirectly with ISG15 through binding to RNF213. Taken together, our Virotrap screens identified RNF213 as a novel, robust non-covalent interaction partner of ISG15 along with a number of VCP and proteasome-associated proteins, all requiring both the N- and C-terminal domains of ISG15 to interact.

### RNF213 binds ISG15 but not ubiquitin or SUMO

To confirm the interaction and specificity between ISG15 and RNF213 through an alternative approach, we first carried out classic immunoprecipitation-MS experiments in IFN-I treated HEK293T and HeLa cells expressing HA-tagged-ISG15AA. In line with previous reports ^38–40^, HA-pull down of ISG15 did not reveal RNF213 nor any other significant interaction partners (Supplementary Fig. 3A-C). This suggested that affinity between a single ISG15 molecule and RNF213 is too weak to detect, similar to ubiquitin or SUMO molecule to their respective binding proteins ^51,52^. Hence, we applied a Glutathione-S-Transferase (GST) pull down assay in which we coated glutathione beads with GST-ISG15 and then used these ISG15- decorated beads to search for ISG15 binding proteins in a lysate of HEK293T, HeLa and THP-1 cells treated with IFN-I. GST-coated beads were used as control and interacting proteins were identified by mass spectrometry. Interestingly, this approach yielded RNF213 as a strongly enriched partner of ISG15 along with several other interactors (Fig. 1D-F), however, only RNF213 was identified in both the GST enrichment and the Virotrap experiments (Supplementary Fig. 3D). Among the other interactors, we detected UBE1L (UBA7) and HDAC6 which are known ISG15 binders in THP-1 and HEK293T cells, respectively ^1,34^. To further confirm the specificity of the ISG15/RNF213 interaction, we next combined the pull down of GST-ISG15 with western blotting. In these experiments, a lysate of HEK293T cells expressing FLAG-RNF213 was mixed with glutathione beads bound to GST-ISG15, GST-SUMO or GST-ubiquitin. While RNF213 clearly binds to GST-ISG15, only background binding could be observed to GST-SUMO and GST-ubiquitin (Fig. 1G). As a positive control, we confirmed binding of RNF4 and RNF31, known binding partners of GST-SUMO and GST-ubiquitin, respectively ^51,53^ (Fig. 1H-I). Together, these data show that RNF213 specifically binds ISG15, rather than ubiquitin or SUMO.

### RNF213 binds ISGylated proteins on lipid droplets

RNF213, also known as mysterin, is a multidomain megaprotein of unknown function. Polymorphisms in *RNF213* predispose human patients to Moyamoya disease (MMD), a rare vascular brain disease leading to stroke, but the underlying molecular mechanisms of this disease remain elusive ^54^. The most abundant RNF213 isoform is 5,207 amino acids long and is comprised of two adjacent AAA+ ATPase modules with a RING domain ^55^. RNF213 is expressed in most cells and tissues ^56^, but levels can be upregulated by immune and inflammatory signaling ^55^. AAA+ proteins typically assemble into hexameric toroidal complexes that generate mechanical force through ATP binding/hydrolysis cycles. In the case of RNF213, complex formation is dynamic and driven by ATP binding while ATP hydrolysis mediates disassembly ^55^. Interestingly, a recent microscopy study showed co-localization of RNF213 and lipid droplets (LDs), suggesting a model in which monomeric RNF213 is directly recruited from the cytosol and ATP binding drives its oligomerization on the surface of lipid droplets ^57^. This model, together with the observation that ISG15 binds RNF213 in Virotrap and GST pull down experiments - which both present multiple ISG15 bait molecules in close proximity on a concave or convex surface, respectively (Fig. 1A) - led us to hypothesize that RNF213 oligomerizes on LDs into a binding platform for ISG15 and potentially multiply ISGylated proteins.

To test this hypothesis, we first verified whether RNF213 can bind ISGylated proteins. To this end, we pulled down FLAG-RNF213 from lysates of HEK293T cells expressing HA-tagged ISG15 or ISG15AA and its conjugation machinery to induce ISGylation. Western blotting against the HA-tag revealed a smear of ISGylated proteins after FLAG-RNF213 pulldown, but not with FLAG-eGFP pulldown or when ISG15AA was expressed (Fig. 2A). We repeated the experiment with eGFP-tagged RNF213 (Supplementary Fig. 4A) and in both cases did not observe any free ISG15 or ISG15AA associated with RNF213, indicating that in cells RNF213 indeed preferentially senses and binds ISGylated proteins rather than free ISG15. In order to determine whether the ISGylated proteins detected were instead partially degraded RNF213 we included a 1% SDS wash, which reduced the ISGylated proteins (Supplementary Fig. 4B) and even demonstrated that RNF213 could bind ISGylated proteins in trans which originated from a different cell lysate (Supplementary Fig. 4C-D). These data indicate that RNF213 does bind ISGylated proteins. Finally, we also showed that this binding was ISG15 specific, since ubiquitinated proteins were not enriched (Supplementary Fig. 4E), in line with the results from the GST-pull down (Fig. 1G).

**Figure 2.**
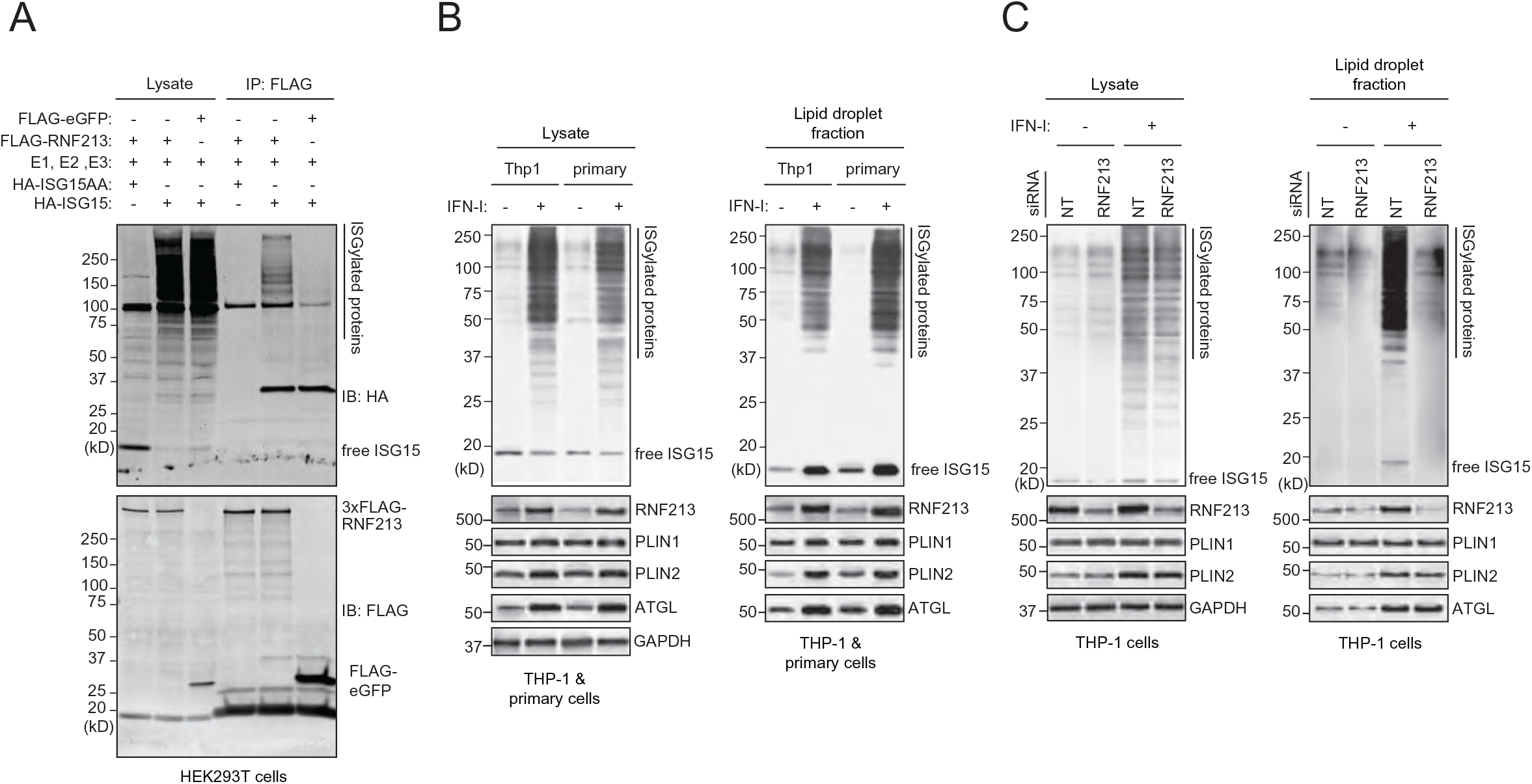
RNF213 binds ISGylated proteins on lipid droplets. **(A)** FLAG immunoprecipitation was performed from lysates of HEK293T cells expressing FLAG-RNF213 or FLAG-eGFP in combination with HA-ISG15(AA) and the ISGylation machinery (E1, E2, E3). A smear of ISGylated co-immunoprecipitated proteins was detected with FLAG-RNF213, but not with FLAG-eGFP or when non-conjugatable HA-ISG15AA was used. **(B)**. THP-1 or primary human monocytes (CD14+) cells were cultured in the presence of 10 mM BSA-conjugated oleic acid and either treated with 10 ng/mL interferon-β for 8 h or left untreated. Lipid droplets (LDs)-enriched fractions were isolated by ultracentrifugation floatation assay on a sucrose step-gradient. Immunoblots against RNF213 and ISG15 revealed an interferon-induced upregulation of both proteins on LDs and a smear of ISGylated proteins associated with LDs. Immunoblots against PLIN1, PLIN2, ATGL and GAPDH confirmed LD isolation and equal protein loading in the lysate and LD-enriched fraction (1/20^th^ of the lysate and all of the LD-enriched material was loaded). **(C)** Similarly, LDs were isolated from THP1 cells after knockdown of RNF213 by siRNA treatment for 48h or using a non-targeting scrambled siRNA (siScramble) as control. Immunoblotting against ISG15 revealed a smear of ISGylated proteins associated with LDs only when RNF213 was present. Immunoblotting against RNF213 confirmed knockdown of RNF213, while PLIN1, PLIN2, ATGL and GAPDH validated LD isolation and equal protein loading in the lysate and the LD- enriched fraction (1/20^th^ of the lysate and all of the LD-enriched material was loaded).

We next tested whether the interaction between RNF213 and ISGylated proteins occurs on LDs. In HeLa cells treated with IFN-I we observed occasional co-localization of overexpressed eGFP-RNF213 with endogenous ISG15 by fluorescence microscopy, however, ubiquitous intracellular distribution of free ISG15 prevented reliable quantitation of these events (data not shown). We therefore verified this question further biochemically using macrophages, a cell type which is more relevant for innate immune signaling. We isolated LDs from THP-1 cells and primary human monocytes by flotation on a sucrose gradient ^58^. Western blotting for RNF213 and LD marker proteins such as PLIN1 and PLIN2 confirmed that in THP-1 cells the majority of RNF213 is associated with LDs, while a minor fraction is in the cytosol (Supplementary Fig. 5A). Upon IFN-I treatment, the fraction of RNF213 associated with LDs further increased, along with a striking appearance of a smear of ISGylated proteins, an observation that was corroborated in primary human monocytes (Fig. 2B). Finally, to show that ISGylated proteins associate with LDs via RNF213, we repeated the experiment in THP-1 cells in which we reduced the expression of RNF213 by siRNA. As expected, knockdown of RNF213 markedly reduced association of ISGylated proteins with LDs and this occurred without any apparent loss of lipid droplet stability as indicated by unchanged levels of LD markers PLIN1, PLIN2 and ATGL (Fig. 2C). Furthermore, in bone marrow-derived macrophages (BMDMs) isolated from RNF213 knockout mice ^59^ we found that the number and size of lipid droplets was similar to WT BMDMs (Supplementary Fig. 5B-C), indicating that in macrophages RNF213 is not essential for lipid droplet stability, in contrast to other cell types ^57^. Together, these data show that RNF213 binds ISGylated proteins and that RNF213 recruits ISGylated proteins to LDs.

### IFN-I induces RNF213 ISGylation and oligomerization on lipid droplets

We subsequently investigated how oligomerization of RNF213 on LDs is regulated. Since both ISG15 and RNF213 are IFN-I induced genes (Supplementary Fig. 6A) ^60^, it seemed plausible that multimerization of RNF213 into a sensing platform for ISGylated proteins could be mediated by interferon signaling. We therefore evaluated the effect of IFN-I on the distribution of endogenous RNF213 between the soluble and membrane-associated fractions in HeLa cells and found that the level of membrane-associated RNF213 slightly increased upon IFN-I treatment (Supplementary Fig. 6B). To further confirm RNF213 oligomerization in a relevant model, we separated lysates from control and IFN-I treated THP-1 cells by sedimentation over continuous glycerol gradients. Western blotting of the collected fractions clearly showed that IFN-I treatment gave rise to a pool of higher molecular weight RNF213 in fractions 14 to 20, as expected after oligomerization (Fig. 3A). Interestingly, this pool of slow migrating RNF213 was marked by the presence of higher molecular weight bands on the blots, most likely representing modified forms of RNF213. Since we recently identified RNF213 as the most ISGylated protein with twenty-two ISGylation sites in the liver of bacterially infected mice ^61^, we tested whether oligomeric RNF213 was ISGylated. Immunoprecipitating RNF213 from each fraction followed by immunoblotting for ISG15 showed that this was indeed the case, while a ubiquitin-specific signal was only detected in the low-density fractions (Fig. 3A). The ISG15 blot revealed two major groups of ISGylated RNF213 with one group of bands running at virtually the same height as monomeric RNF213 and a second, hyper-ISGylated group of bands running well above (Fig. 3A). Knockdown of Ube1L, the E1 activating enzyme for ISG15, strongly reduced the intensity of both groups of ISGylated RNF213 while it increased the intensity of monomeric RNF213, indicating that ISGylation of RNF213 is required for its oligomerization (Fig. 3B and Supplementary Fig. 6C). Finally, we repeated the glycerol sedimentation experiment after LD isolation and showed that the oligomeric, ISGylated forms of RNF213 are associated with LDs only after IFN-I treatment (Fig. 3C). Together, these results show that IFN-I induces the oligomerization of RNF213 on LDs and that this process requires ISG15 modification of RNF213 itself.

**Figure 3.**
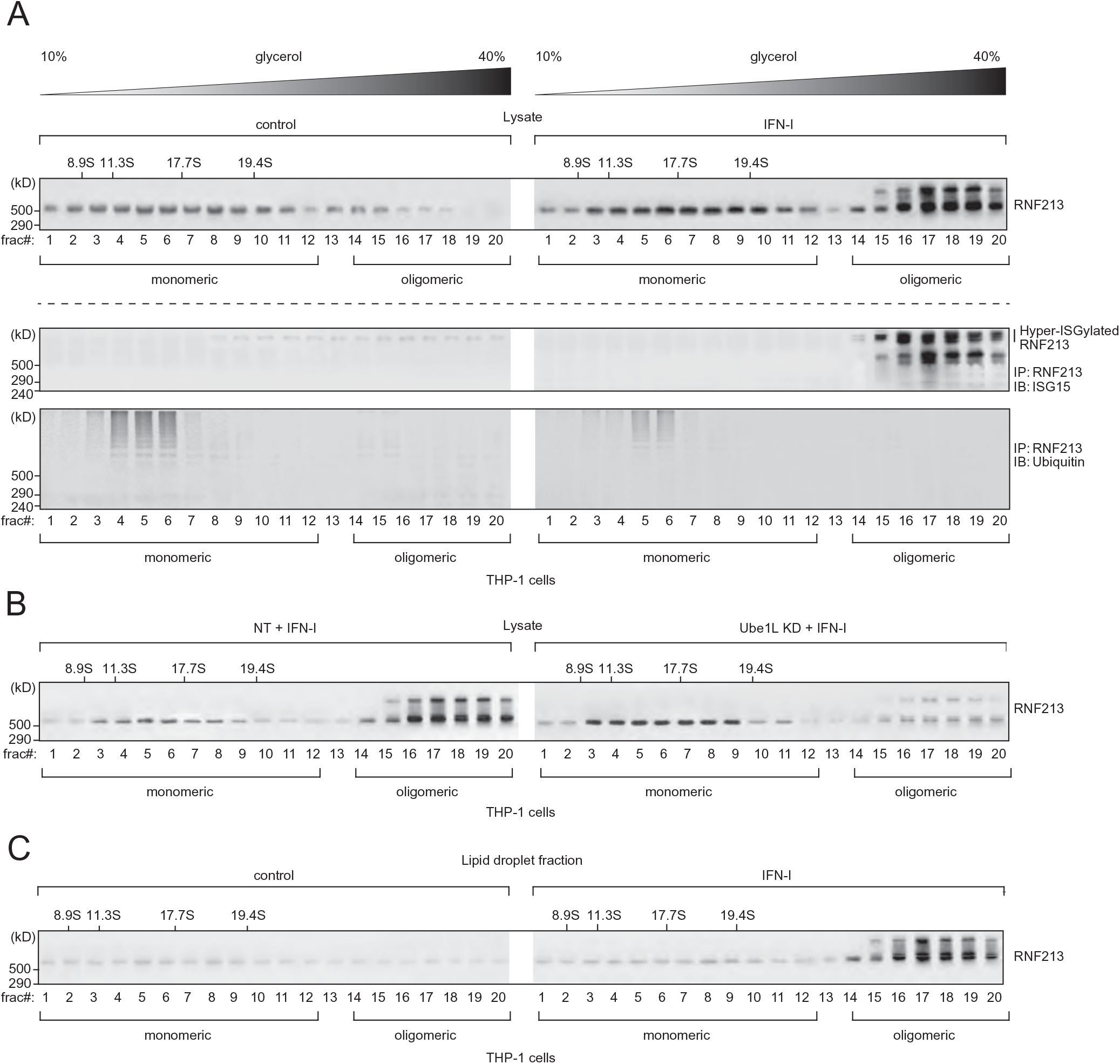
Type-I interferon induces RNF213 ISGylation and oligomerization on lipid droplets. **(A)** THP-1 cells were treated with 10 ng/mL interferon-β for 8 h or left untreated. Lysates were separated by density gradient ultracentrifugation on glycerol gradients (10 – 40% (v/v), Svedberg constants of the standard markers are indicated above the blots) to isolate the monomeric versus oligomeric form of RNF213. Twenty fractions for each sample were collected, concentrated by TCA precipitation and analyzed by immunoblotting against RNF213, showing the presence of oligomer RNF213 in fraction 14 to 20 upon interferon treatment (upper panel). Alternatively, RNF213 was immunoprecipitated (IP) from each fraction, first desalted over Amicon columns. Immunoprecipitated material was eluted into loading buffer and analyzed by immunoblotting against ISG15 and ubiquitin, showing (hyper)ISGylation of oligomer RNF213 upon interferon treatment (lower panel). (**B**) The monomeric and oligomeric forms of RNF213 were separated by density gradient ultracentrifugation after interferon-β treatment as in (A) and knockdown of Ube1L by siRNA treatment for 48 h, using a non-targeting scrambled siRNA (NT) as control. Knockdown of Ube1L strongly reduced RNF213 oligomerization upon interferon treatment. **(C)** LDs were isolated by ultracentrifugation floatation assay on a sucrose step-gradient and associated proteins were further separated by density gradient ultracentrifugation to isolate the monomeric and oligomeric form of RNF213 after interferon-β treatment as in (A). Fractions were concentrated by TCA precipitation and analyzed by immunoblotting against RNF213, showing association of oligomeric RNF213 with LDs upon interferon treatment.

### RNF213 exerts broad antimicrobial activity *in vitro*

The discovery of RNF213 as a sensor for ISGylated proteins made us hypothesize that RNF213 could also have antiviral properties, similar to ISG15. To investigate this, we infected cells with reduced or enhanced expression levels of RNF213 with different viral pathogens. We first infected HeLa cells with herpes simplex virus 1 (HSV-1), a widespread human pathogen that can cause cold sours and genital herpes that is ISG15-sensitive ^62^. We used a GFP-expressing HSV-1 strain to monitor infection levels up to three days post infection by fluorescence intensity. Knockdown of RNF213 led to significantly higher infection levels compared to scrambled siRNA control, both in the presence or absence of IFN-I (Fig. 4A-C). Conversely, overexpression of FLAG-RNF213 significantly lowered HSV-1 infection levels, although not to the same extent as overexpressed MXB, which was recently reported as a Herpesvirus restriction factor (Fig. 4D-E)^63^. We also determined the possible involvement of RNF213 in the control of replication of human respiratory syncytial virus (RSV), an important human respiratory pathogen that is susceptible to an ISG15- dependent antiviral effect ^10^. For these experiments, A549 cells were infected with RSV-A2 at a low multiplicity of infection and incubated for six days to allow multiple rounds of infection (Supplementary Fig. 7A). Knockdown of RNF213 significantly increased the viral load in the cell culture supernatants five days post-infection compared to scrambled siRNA control, an effect that disappeared at day six (Fig. 4F). In addition, when the cells were treated with IFN-I, significantly higher viral loads were observed in siRNF213-treated cells compared to siScramble-treated cells at day five and six after infection (Fig. 4G). We also assessed the effect of RNF213 knock down on RSV replication in a plaque assay because the plaque size is a measure for the extent of cell-to-cell spread of the virus. On day six after RSV infection, we observed significantly larger plaques upon knockdown of RNF213 both with and without IFN-I, suggesting also increased cell-to-cell spread of RSV under these conditions (Fig. 4H-I). As a third ISG15- sensitive viral pathogen we tested CVB3, a Picornavirus that can lead to cardiomyopathy and that is known to be ISG15-sensitive ^11,12^ (Supplementary Fig. 7B). Again, knockdown of RNF213 significantly increased the replication of CVB3 as indicated by higher viral genome (Fig. 4J) and enhanced viral protein expression levels (Fig. 4K) as well as by the elevated formation of infectious viral particles from cells (Fig. 4L).

**Figure 4.**
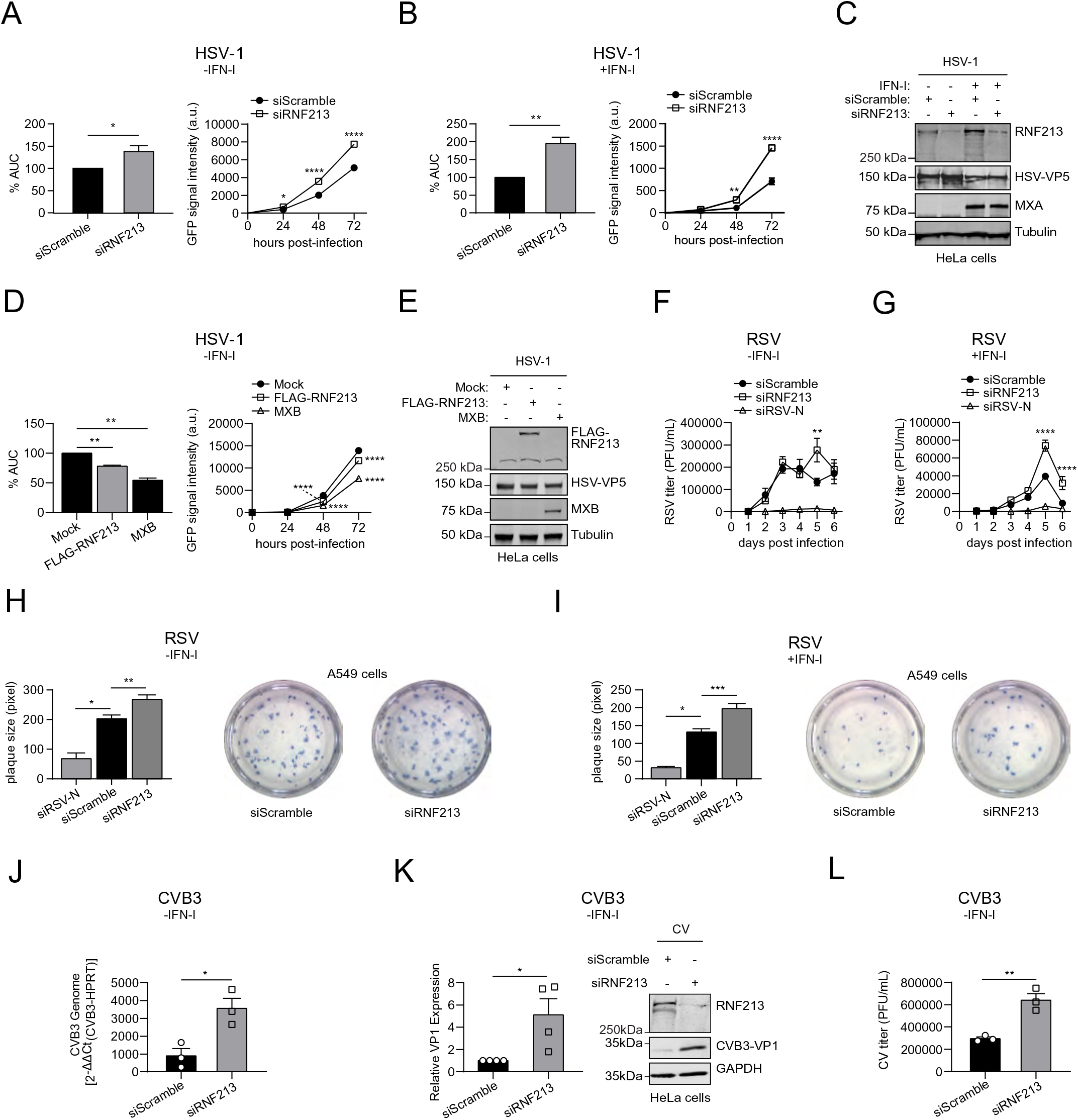
RNF213 counteracts herpes simplex, respiratory syncytial and coxsackie virus infection. **(A)** HeLa cells were infected up to 72 h with eGFP-expressing recombinant herpes simplex virus 1 (HSV-1) at MOI 0.1. 72 h prior to infection, cells were transfected with a pool of siRNAs targeting RNF213 (siRNF213) or a pool of scrambled siRNAs (siScramble) as control. The viral load was determined by monitoring the GFP signal in each condition every 24 h to generate a viral growth curve (right panel, representative viral growth curve, AVG ± SEM, n=4 replicates, two-tailed unpaired t-test comparing siRNF213 to siScramble, * p < 0.05, ** p < 0.01 and **** p < 0.0001). The area under the curve (AUC) was calculated for each growth curve and the average AUC of three independent experiments is shown relative to the siScramble control (left panel, AVG ± SEM, n=3 independent experiments, two-tailed unpaired t-test, * p < 0.05 and ** p < 0.01). **(B)** HSV-1 infection experiment performed as in (A), except that 16 h prior to infection cells were treated with 1,000 U/mL interferon-α. Knockdown of RNF213 leads to significantly higher HSV-1 infection levels, both in the absence (A) and presence of interferon-α (B). **(C)** Immunoblots against RNF213, HSV-VP5 and MXA with tubulin as loading control confirmed knockdown of RNF213, HSV-1 infection and interferon-α treatment, respectively, in the experiments shown in (A-B). **(D)** HSV-1 infection experiment performed as in (A-B), except that 24 h prior to infection cells were transfected with plasmids encoding 3X-FLAG-RNF213 or MXB or with an empty vector (mock) as control (right panel, representative viral growth curve, AVG ± SEM, n=4 replicates, two-tailed unpaired t-test comparing siRNF213 to siScramble, * p < 0.05, ** p < 0.01 and **** p < 0.0001). The average AUC of two independent experiments is shown relative to the mock control (left panel, AVG ± SEM, n=2 independent experiments, two-tailed unpaired t-test, * p < 0.05 and ** p < 0.01). Overexpression of RNF213 leads to significantly lower HSV-1 infection levels, as also observed for MXB as positive control. **(E)** Immunoblots against FLAG, HSV-VP5 and MXB with tubulin as loading control confirmed HSV-1 infection and expression of FLAG-RNF213 and MXB in the experiments shown in (D). **(F)** A549 cells were infected with RSV-A2 for up to six days at MOI 0.02. 48 h prior to infection, cells were transfected with a pool of siRNAs targeting RNF213 (siRNF213), a single siRNA targeting the RSV-nucleoprotein (RSV-N) as positive control or a pool of scrambled siRNAs (siScramble) as negative control. The viral titer was determined by counting plaque-forming units (PFUs) after serial dilution (AVG ± SEM, n=3 replicates, two-tailed unpaired t-test compared siRNF213 to siScramble, ** p < 0.01, and **** p < 0.0001). **(G)** RSV infection experiment performed as in (F), except that 42 h prior to infection cells were treated with 10 ng/mL interferon-β. Knockdown of RNF213 leads to significantly higher RSV titers, both in the absence (F) and presence of interferon-β (G). **(H)** A549 cells were infected with RSV-A2 at MOI 0.005 in combination with knockdown of RNF213 and RSV-N as described in (F). Six days post infection, a plaque assay was performed and plaque sizes were quantified in pixels with Fiji (left panel, AVG ± SEM, Mann– Whitney test, * p < 0.05, ** p < 0.01 and *** p < 0.001, siRSV-N n=8, siScramble n=267 and siRNF213 n=183). Representative images showing plaques of the siRNF213 and siScramble treated cells are shown in the right panel. **(I)** RSV infection experiment performed as in (H), except that 42 h prior to infection cells were treated with 10 ng/mL interferon-β (left panel, siRSV-N n=4, siScramble n=123 and siRNF213 n=99). Knockdown of RNF213 leads to significantly larger RSV plaques, both in the absence (H) and presence of interferon-β (I). **(J)** HeLa cells were infected with coxsackievirus B3 (CVB3) at MOI 0.01. 24 h prior to infection, cells were transfected with a pool of siRNAs targeting RNF213 (siRNF213) or a pool of scrambled siRNAs (siScramble) as control. 24h post infection, the intracellular viral RNA load was determined by qRT-PCR (AVG ± SEM, n=3 independent experiments, two-tailed unpaired t-test, * p < 0.05). **(K)** From the same experiment as in (J), the intracellular viral protein load was determined by immunoblotting against VP1 and the intensity of the VP1 band is shown relative to the siScramble control (left panel, AVG ± SEM, n=4 independent experiments, one-tailed t-test, * p < 0.05). A representative immunoblot for the quantification of VP1 is shown in the right panel. **(L)** From the same experiment as in (J), the viral titer was determined by counting PFUs after serial dilution (AVG ± SEM, n=3 replicates, two- tailed unpaired t-test, ** p < 0.01). Knockdown of RNF213 leads to a significant increase in CVB3 infection as measured by higher viral genome (J), protein (K) and titer (L).

We subsequently wondered whether RNF213 could be broadly antimicrobial and also target bacterial pathogens. Thus, we infected HeLa cells with *Listeria monocytogenes*, a facultative intracellular bacterium that was recently reported to be counteracted by ISG15 ^6^ (Supplementary Fig. 7C). When we knocked down RNF213 with a pool of siRNAs, we measured a 30% increase in intracellular bacteria compared to a pool of control siRNA, an increase that was similar to what we observed by knockdown of ISG15 (Fig. 5A). Individual siRNAs against RNF213 also augmented the bacterial load (Supplementary Fig. 7D). Concordantly, overexpression of FLAG-RNF213 reduced the bacterial load with 85% relative to mock transfected cells, a reduction that was similar to overexpression of HA-ISG15 (Fig. 5B). Importantly, the protective effect of overexpressed FLAG-RNF213 was lost in ISG15 knockout cells (Fig. 5C), indicating that the antimicrobial activity of RNF213 requires ISG15. In contrast, overexpressed HA-ISG15 still counteracted *Listeria* in RNF213 knockout (Fig. 5D) or knockdown cells (Supplementary Fig. 7E), meaning that ISG15 also harbors antimicrobial activity independent of RNF213. Taken together, we showed that increasing the expression levels of RNF213 led to lower infection levels of HSV-1 and *L. monocytogenes* in cultured cells, while reducing the levels of RNF213 promoted *in vitro* infection with HSV-1, RSV, CVB3 and *L. monocytogenes*. These results show that RNF213 plays an important role in the innate cellular immune response, indicating a functional link between ISG15 and RNF213.

**Figure 5.**
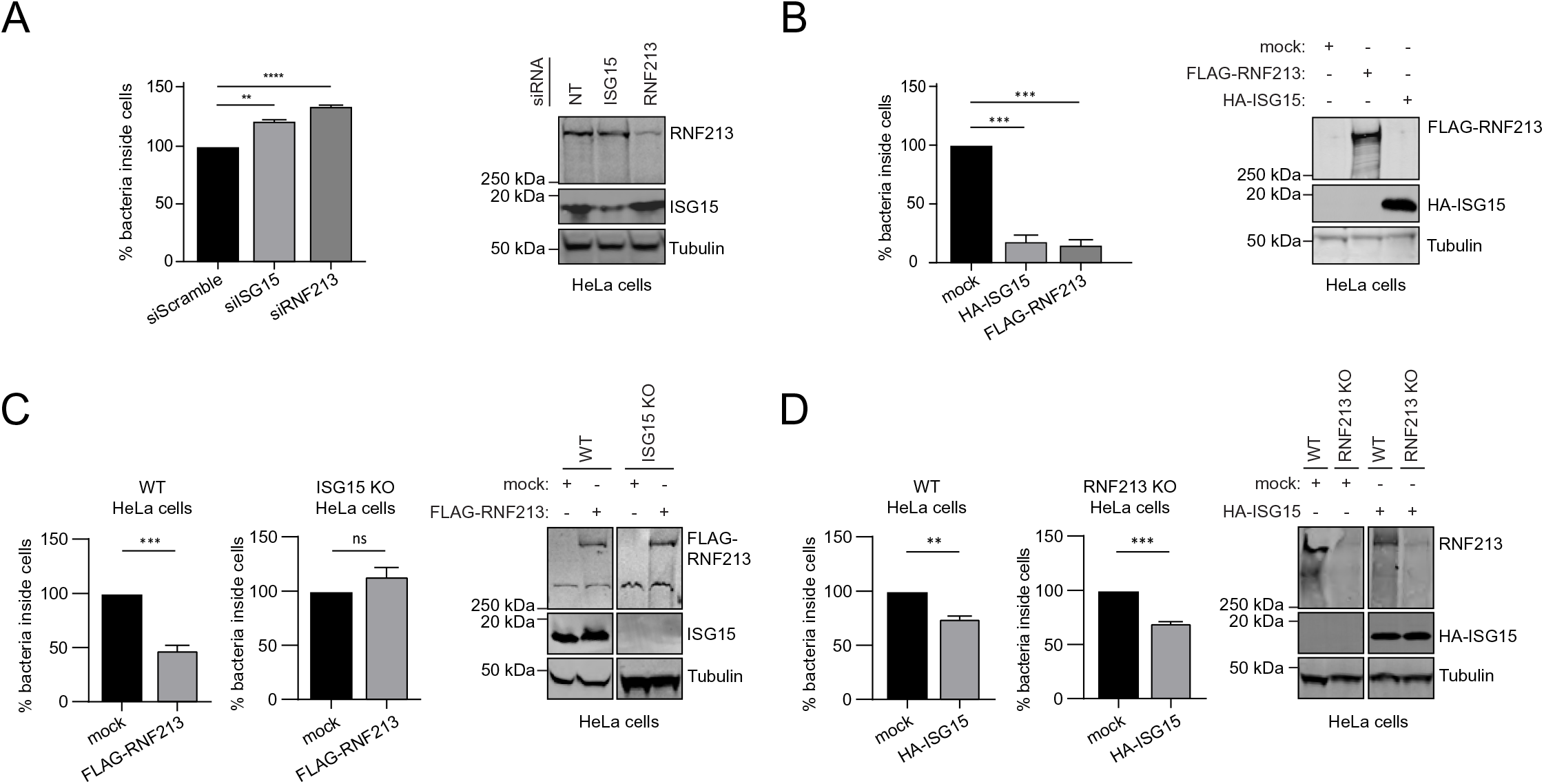
RNF213 counteracts *Listeria* infection. **(A)** HeLa cells were infected with *Listeria monocytogenes* EGD for 16 h at a multiplicity of infection (MOI) of 25. 24 h prior to infection, cells were transfected with a pool of siRNAs targeting ISG15 (siISG15), RNF213 (siRNF213) or a pool of scrambled siRNAs as control. Intracellular *Listeria* were quantified after serial dilution by counting colony-forming units (CFUs) in a gentamycin assay. The percentage of intracellular bacteria relative to siScramble-transfected cells is shown (AVG ± SEM, n=3 independent experiments, two-tailed unpaired t-test, ** p < 0.01, **** p < 0.0001) (left panel). Immunoblots against RNF213 and ISG15 with tubulin as loading control confirmed knockdown of both proteins (right panel). **(B)** HeLa cells were infected with *Listeria* for 4 h at MOI 25. 24 h prior to infection, HeLa cells were transfected with plasmids encoding 3xFLAG-RNF213 or HA-ISG15 or with an empty vector (mock) as control. Intracellular *Listeria* were quantified as in (A) and the percentage of intracellular bacteria relative to mock plasmid-transfected cells is shown (AVG ± SEM, n=3 independent experiments, two-tailed unpaired t-test, *** p < 0.001) (left panel). Immunoblots against FLAG and HA with tubulin as loading control confirmed expression of both proteins (right panel). **(C)** Wild-type (WT) or ISG15 knockout HeLa cells were infected with *Listeria* for 16 h at MOI 25 after transfection of FLAG-RNF213 as in (B). Intracellular *Listeria* were quantified as in (A) and the percentage of intracellular bacteria relative to mock plasmid-transfected cells is shown (AVG ± SEM, n=3 independent experiments, two-tailed unpaired t-test, *** p < 0.001) (left panel). Immunoblots against FLAG and ISG15 with tubulin as loading control confirmed expression of FLAG-RNF213 and the absence of ISG15 in the KO cells (right panel). **(D)** Wild- type (WT) or RNF213 knockout HeLa cells were infected with *Listeria* for 16 h at MOI 25 after transfection of HA-ISG15 as in (B). Intracellular *Listeria* were quantified as in (A) and the percentage of intracellular bacteria relative to mock plasmid-transfected cells is shown (AVG ± SEM, n=3 independent experiments, two-tailed unpaired t-test, ** p < 0.01, *** p < 0.001) (left panel). Immunoblots against RNF213 and HA with tubulin as loading control confirmed expression of HA-ISG15 and the absence of RNF213 in the KO cells (right panel).

### RNF213 decorates intracellular *Listeria monocytogenes* and is profoundly antibacterial *in vivo*

Given the protective function of RNF213 against *Listeria*, we wondered if RNF213 adopts a specific subcellular localization during infection. To this end, we transfected HeLa cells with eGFP-RNF213. Co-staining for LDs followed by fluorescence microscopy revealed that RNF213 co-localized with the majority (∼70%) of LDs, both in uninfected and *Listeria*-infected cells (Fig. 6A-B). Intriguingly, in the latter condition RNF213 also co-localized with a subset of intracellular *L. monocytogenes*, decorating the bacterial surface. To quantify this phenomenon we repeated the experiment with mCherry-expressing *Listeria* and found that on average approximately 40% of intracellular Listeria co-colocalized with RNF213 (Fig. 6C-D). Thus, our data show that during infection with *L. monocytogenes*, in addition to LDs, RNF213 localizes to a subset of intracellular bacteria.

**Figure 6.**
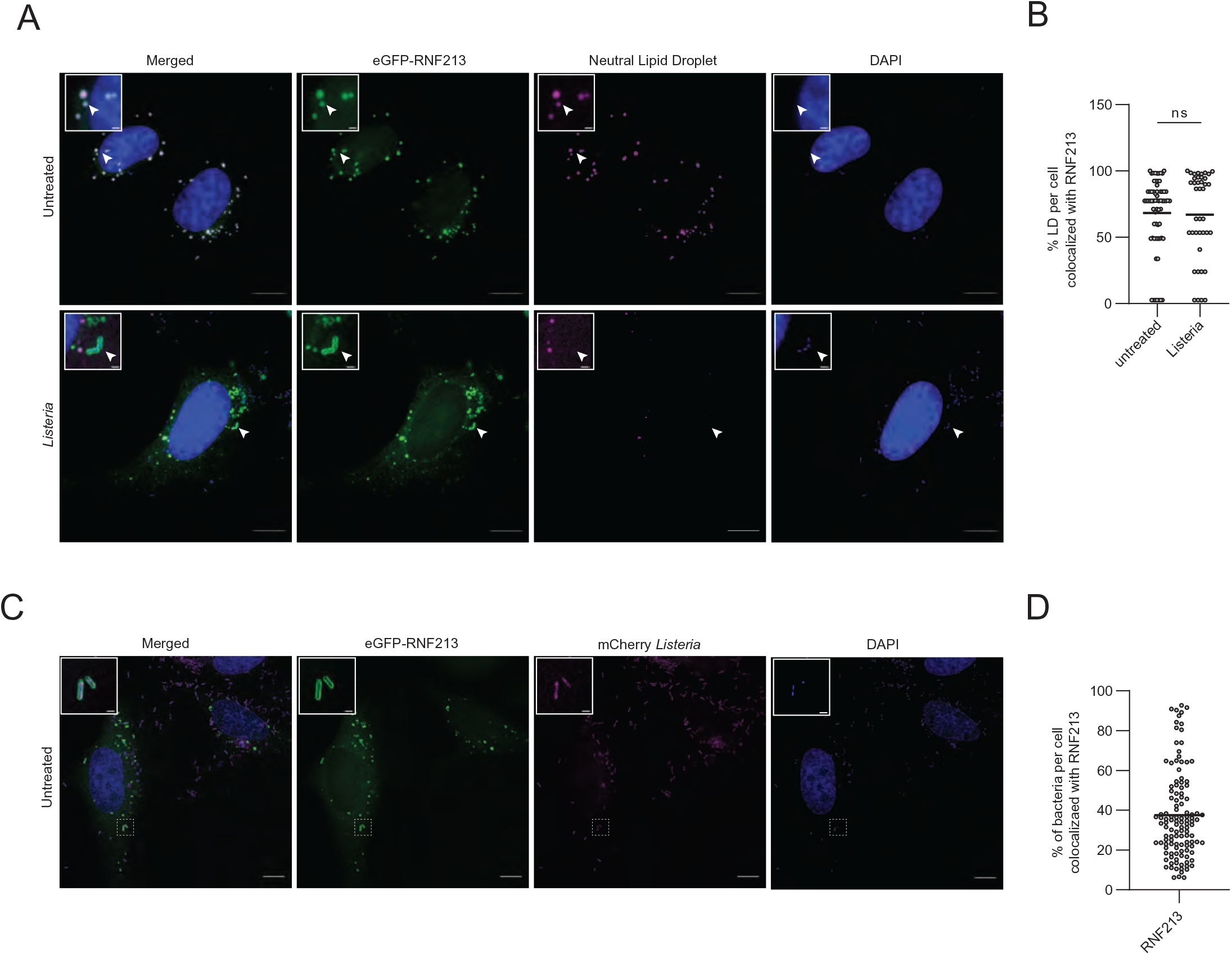
RNF213 decorates intracellular *Listeria*. **(A)** Representative images of HeLa cells transfected with eGFP-RNF213 and counterstained for lipid droplets (LDs). Following transfection cells were left untreated or infected with *Listeria monocytogenes* EGD for 24 h. Scale bars in the pictures and insets are respectively 10 microns and 1 micron. White arrows indicate co-localization between RNF213 and lipid droplets or instances of RNF213 co-localization with intracellular bacteria. **(B)** LDs in uninfected (n=66 cells) and *Listeria*-infected cells (n=40 cells) from (A) were quantified with Fiji and the percentage of LDs per cell that co-localized with RNF213 was calculated, showing no difference in co-localization between uninfected and *Listeria*-infected cells (two-tailed unpaired t-test, AVG uninfected = 68.29%, AVG Listeria-infected = 67.14%). **(C)** Representative images of HeLa cells transfected with eGFP-RNF213 and infected for 18 hours with *Listeria monocytogenes* EGD stably expressing mCherry. Scale bars in the pictures and insets are respectively 10 microns and 0.5 micron. **(D)** Intracellular *Listeria* from (C) were quantified with Imaris 9.6 and the percentage of *Listeria* that was decorated by eGFP-RNF213 was calculated for each field by mapping the cell surface, enumerating intracellular bacteria and quantifying bacteria that colocalized with GFP-RNF213 (defined as bacteria within 0.5 µm of RNF213, see methods for more detail). At least 200 cells were counted per experiment, and data were compiled from 3 independent experiments indicating that on average 37.44% of *Listeria* was decorated by RNF213.

In order to test whether this localization has functional consequences *in vivo*, we deleted RNF213 in mice using CRISPR/Cas9 targeting of mouse *RNF213* exon 28 as previously described ^59^ (Supplementary Fig. 8A). At 24, 48 and 72 hours post infection with *L. monocytogenes* we observed a dramatic increase in bacterial burden in the spleen of *rnf213*^-/-^ animals compared to WT littermate controls which increased over time (Fig. 7A). At 72 hours, a significant increase was also detected in the liver (Fig. 7B) and this observation was confirmed in two additional independent experiments (Supplementary Fig. 8B-E). This profound difference in bacterial load, particularly in the spleen, reveals the central importance of RNF213 in immune cell function and intracellular bacterial clearance. Future work will assess which cell type is unable to control infection and whether RNF213 deletion also affects adaptive next to innate responses. This is of particular relevance to understanding the role of inflammation or prior infection in human patients with inborne errors in RNF213.

**Figure 7.**
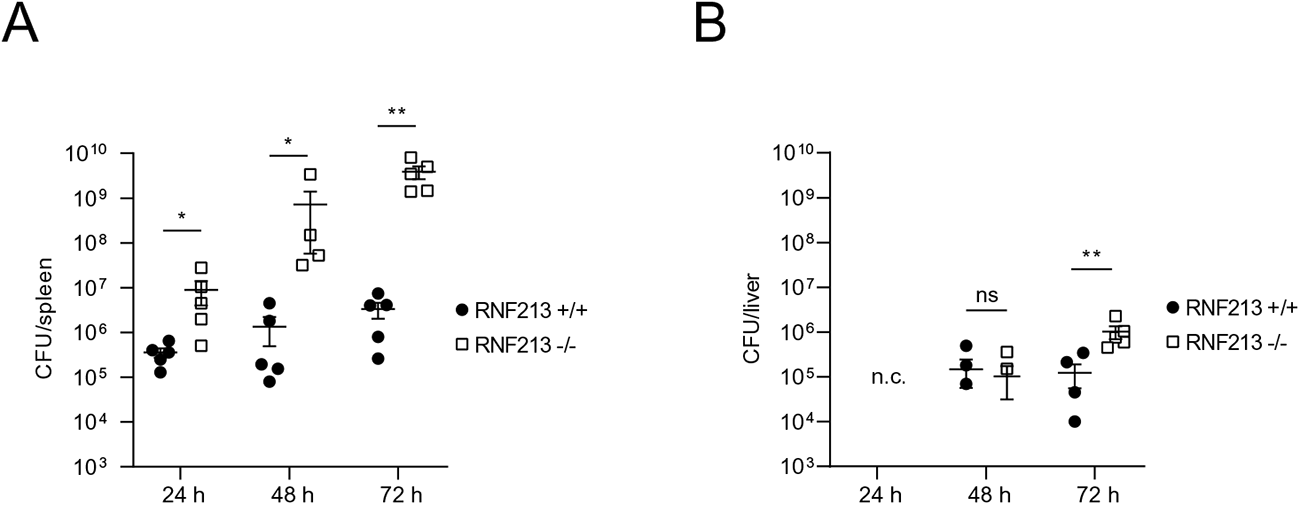
RNF213 counteracts *Listeria* infection *in vivo*. **(A-B)** RNF213 -/- and RNF213 +/+ mice were infected intravenously with 5 × 10^5^ *Listeria monocytogenes* EGD. Spleen (A) and liver (B) were isolated following 24 h (n=5 for both genotypes), 48 h (n= 4 for RNF213 -/- and n=5 for RNF213 +/+) and 72 h of infection (n=5 for both genotypes). Colony forming Units (CFUs) per organ were counted by serial dilution and replating; dots and squares depict individual animals (AVG ± SEM, Mann–Whitney test, * p < 0.05 and ** p < 0.01, five data points fell below detection limit in the liver). RNF213 -/- mice are dramatically more susceptible to *Listeria* as evidenced by significantly higher CFUs at all three time points in the spleen and at 72 h in the liver.

### The RNF213 E3 module is required for its antimicrobial activity

While the present study was under review, Ahel et al. published the structure of monomeric RNF213 ^64^ showing that RNF213 does not depend on its RING domain to function as an E3 ubiquitin ligase, but instead classifies as a new type of E3 ligase employing a Cys-containing motif to promote ubiquitin transfer via transthiolation. While the active site cysteine residue remains to be determined and a ΔRING mutant still exhibits ubiquitination activity ^64^, we sought to use this newly available structural data to test whether the E3 ligase activity of RNF213 is required for its antimicrobial function. We therefore deleted the C-terminal 1210 amino acids of RNF213 comprising the large E3 module and small C-terminal domain to generate a FLAG-RNF213ΔC construct that should be devoid of any ubiquitination activity. GST pull down showed that RNF213ΔC was still capable of binding ISG15, while the complementary C-terminal fragment was not (Fig. 8A). Similarly, FLAG-RNF213ΔC was capable of pulling down a smear of ISGylated proteins to the same extent as full length FLAG-RNF213 (Fig. 8B). However, interestingly overexpression of RNF213ΔC no longer retained its notable antimicrobial effects when overexpressed as it could not reduce the amount of intracellular *Listeria* in infected HeLa cells (Fig. 8C). Furthermore, fluorescence microscopy confirmed that eGFP-RNF213ΔC did no longer co-localize with intracellular *Listeria* (Fig. 8D) nor with LDs (Supplementary Fig. 9A-B). Taken together, these results indicate that the E3 module of RNF213, and likely its ubiquitination activity, is required for its bacterial targeting and antimicrobial activity.

**Figure 8.**
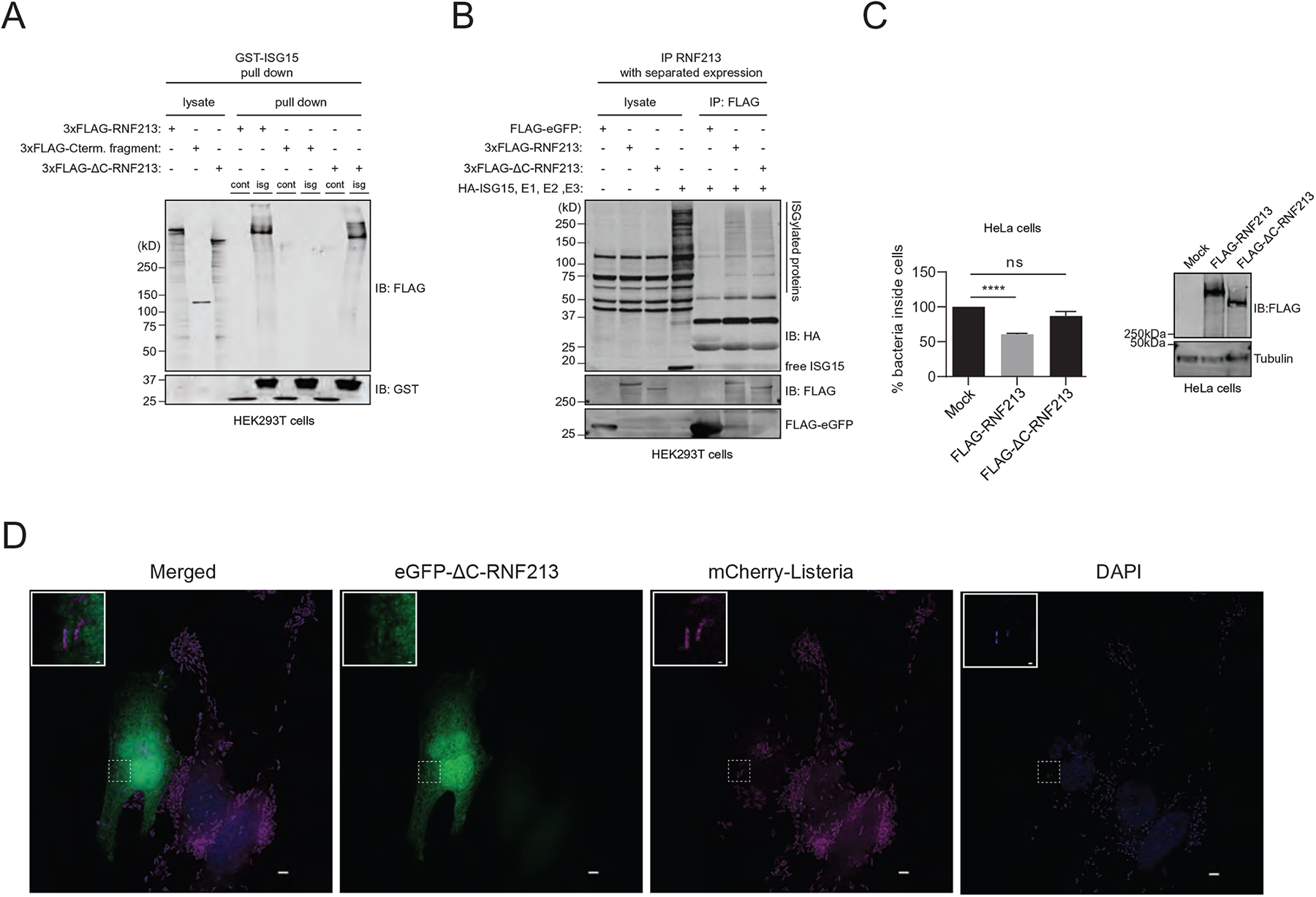
The RNF213 E3 module is required for its antimicrobial activity. **(A)** GST pull down with ISG15 followed by immunoblotting shows that 3xFLAG-RNF213ΔC binds to ISG15 similar to full length 3xFLAG-RNF213. Beads coated with GST-ISG15 were mixed with a lysate of HEK293T cells expressing full length 3xFLAG-RNF213, 3xFLAG-RNF213ΔC or the complementary C-terminal fragment of RNF213 and bound proteins were analyzed by immunoblot against FLAG and GST. (**B**) FLAG immunoprecipitation was performed from lysates of HEK293T cells expressing 3xFLAG- RNF213, 3xFLAG-RNF213ΔC or FLAG-eGFP as control. After immunoprecipitation, beads were mixed with a lysate of HEK293T cells expressing HA-ISG15 and the ISGylation machinery (E1, E2, E3). 3xFLAG-RNF213ΔC was capable of pulling down ISGylated proteins similar to 3xFLAG-RNF213, while FLAG-eGFP was not. **(C)** HeLa cells were infected with *Listeria monocytogenes* EGD for 4 h at MOI 25. 24 h prior to infection, HeLa cells were transfected with plasmids encoding 3xFLAG-RNF213 or 3xFLAG- RNF213ΔC or with an empty vector (mock) as control. Intracellular *Listeria* were quantified after serial dilution by counting colony-forming units (CFUs) in a gentamycin assay. The percentage of intracellular bacteria relative to mock plasmid-transfected cells is shown (AVG ± SEM, n=3 independent experiments, two-tailed unpaired t-test, **** p < 0.0001) (left panel). Immunoblots against FLAG with tubulin as loading control confirmed expression of FLAG-RNF213, FLAG-RNF213ΔC (right panel). **(D)** Representative images of HeLa cells transfected with eGFP-RNF213ΔC and infected for 18 hours with *Listeria monocytogenes* EGD stably expressing mCherry. Scale bars in the pictures and insets are respectively 10 microns and 0.5 micron. eGFP-RNF213ΔC showed a diffused cellular staining, not decorating intracellular *Listeria*.

## DISCUSSION

We report the discovery of RNF213 as an intracellular sensor for proteins modified by ISG15, a ubiquitin-like, immunity-related modification. We show that IFN-I signaling induces the oligomerization and translocation of RNF213 on lipid droplets in macrophages and that this process requires ISGylation of RNF213 itself. Furthermore, *in vitro* and *in vivo* infection assays with four different pathogens revealed an as yet undescribed, broad antimicrobial function of RNF213. Thus, our unique Virotrap method revealed a novel functional link between ISG15 and RNF213, as well as an undiscovered role for RNF213 in host defense pathways to both bacterial and viral infections.

We identified RNF213 as an ISG15-binding protein by Virotrap and confirmed the interaction by GST pull down assays, methods that both create a curved surface, either convex or concave, studded with ISG15 molecules which could be a surrogate for multiply modified proteins or larger binding surfaces such as organelles. It was of particular interest, that we identified a large AAA+ ATPase as ISG15 interactors under these conditions, RNF213, which presumably docks on ISG15 surfaces or vice versa. Indeed, our data suggests that oligomeric RNF213 acts as a binding platform for ISGylated proteins on the surface of lipid droplets. In line with this, we demonstrated that ISGylated proteins co-immunoprecipitated with RNF213 and that fewer ISGylated proteins were isolated on lipid droplets without RNF213. It remains unknown what happens to ISGylated proteins after binding to RNF213. As a model, we were inspired by similarities between the interaction of poly-SUMOylated substrate proteins with RNF4, a SUMO-targeted ubiquitin ligase (STUbL) ^51^ that could be an analogous system to RNF213 and ISGylation. In the STUbL pathway, nuclear stresses lead to poly-SUMOylation of protein targets that are further recognized by RNF4. RNF4 then ubiquitinates the poly-SUMOylated proteins, targeting them for degradation by the proteasome. Our model is that RNF213 might recognize multi-ISGylated proteins as the entry point of a novel pathway for further processing of multi-ISGylated proteins at lipid droplets. While RNF4 & poly-SUMOylation are both induced by nuclear stress, RNF213 and multi-ISGylation are induced by IFN-I signaling ^60,65^. Moreover, co-regulation of ISG15 and RNF213 was previously observed in a large-scale analysis of protein quantitative trait loci (pQTL) ^66^, also suggesting that both proteins operate in the same cellular pathway. Our future work will address the fate of ISGylated proteins that bind RNF213. It is particularly intriguing to determine if the E3 module of RNF213 ubiquitinates the bound ISGylated proteins for proteasomal degradation, which could be substantiated by the identification of VCP and proteasome components alongside RNF213 in the ISG15 Virotrap experiments. Alternatively, the ISGylated proteins bound to RNF213 might be degraded via lipophagy ^67^ or targeted for an alternative fate.

RNF213 is a very large, poorly characterized human protein. Recent functional studies highlighted the role of RNF213 in lipid metabolism, mediating lipid droplet formation ^57^ and lipotoxicity ^68^. The latter study also assessed RNF213-dependent ubiquitinome changes and showed that RNF213 was required to activate the nuclear factor κB (NFκB) pathway upon palmitate treatment. This NFκB-inducing function of RNF213 would, however, be negatively regulated by its ubiquitin ligase activity ^69^ and requires further investigation, especially in light of the RNF213 antimicrobial activity reported here. The newly published cryo-EM structure of monomer RNF213 revealed three major structural components comprising an N-terminal stalk, a dynein-like ATPase core and a C-terminal E3 module ^64^. Previous studies showed that the ATPase activity of RNF213 is essential for its multimerization and localization to lipid droplets, however, in contrast to most AAA+ ATPases that form stable hexamers ^55^, multimerization of RNF213 is proposed to be a dynamic process, similar to the dynamic assembly of the microtubule-severing protein katanin ^55,57^. In this model RNF213 exists as a monomer in solution and forms a hexamer on the surface of lipid droplets upon ATP binding ^57^. Contrary to this, the RNF213 structure demonstrated the presence of six AAA units within the monomeric molecule ^64^, putting into question the mechanism of oligomerization and proposed hexameric nature of oligomer RNF213. We here report that oligomerization of RNF213 on lipid droplets is induced by IFN-I and requires ISGylation of RNF213 itself, an observation that fits with our previous identification of RNF213 as the most ISGylated protein in liver of *Listeria*-infected mice ^61^. Various inducers of RNF213 ISGylation could thus potentially facilitate future structural studies on oligomeric RNF213. However, the mechanism of IFN-I induced oligomerization of RNF213 remains an open question. Relevant in this regard is that RNF213 was reported as a substrate of Protein tyrosine phosphatase 1B (PTP1B or PTPN1), controlling non-mitochondrial oxygen consumption in response to hypoxia ^70^. Since PTP1B also affects JAK-STAT signaling ^71,72^, it would be worthwhile testing whether PTPB1 also regulates RNF213 oligomerization in response to interferon.

It is likely that further processing of ISGylated proteins bound to RNF213 is closely linked to the emerging function of LDs beyond passive lipid storage and lipid homeostasis. LDs have a role in the coordination of immune responses as they participate in the production of pro-inflammatory mediators ^73^. LDs have been associated with the IFN-I response and antigen cross presentation ^74^. Moreover, an increasing number of studies show that viral and bacterial pathogens target LDs during infection either for nutritional purposes or as part of an anti-immunity strategy ^73^. Recent work showed that host cells exploit LD targeting by pathogens by loading LDs with antimicrobial proteins as an intracellular first line of defense ^75^. In this study, RNF213 was found to be enriched on LDs isolated from the liver of LPS-treated mice, similar to our results on IFN-treated cells. This is particularly interesting since ISGylation is also induced by LPS ^7^ The localization of RNF213 on the surface of LDs could represent a host-cell strategy for fighting intracellular pathogens that hijack LDs. Given that LDs are also sites of viral assembly, RNF213 could interrupt this process to decrease infectivity in the cell. The broad antimicrobial activity of RNF213 that we observed bolsters this hypothesis.

The localization of RNF213 as a general virus restriction factor on the surface of LDs could be further rationalized in the context of enteroviruses, such as coxsackievirus, which replicate at specialized membranous domains named replication organelles ^76^. However, whether RNF213 also resides in such lipid-rich replication organelles remains to be determined. On the other hand, ISG15 contributes to the IFN-dependent anti-RSV effect, whereas no role for RNF213 has been reported in the control of RSV. In this context, it is important to note that RSV encodes two prominent non-structural proteins that profoundly suppress type I IFN induction and signaling ^77^. The latter is also true for HSV-1, which has evolved multiple strategies to evade the host antiviral response and to establish latent infections, so far without any known involvement of RNF213 ^78^. With all three tested viral pathogens we observed a clear, but subtle antiviral effect of RNF213. Knockdown or overexpression of RNF213 changed viral infection levels two to five fold, in contrast to *Listeria* where we observed up to 100,000 fold differences in bacterial load, especially *in vivo*, indicating a more pronounced antibacterial effect.

Genome-wide siRNA screens in *Listeria*-infected HeLa cells previously ranked RNF213 as a protective host protein *in vitro*, in line with our observations ^79^. Importantly, the protective effect of RNF213 was lost in ISG15 KO cells, indicating that the antimicrobial activity of RNF213 requires ISG15. Inversely, ISG15 was still functional in RNF213 deficient cells, pointing towards other unknown conjugation-dependent or -independent mechanisms by which ISG15 counteracts *Listeria*, as shown for many other pathogens (Perng and Lenschow, 2018). Our infection experiments with *Listeria* in mice which lack RNF213 corroborated the protective effect *in vivo*, showing a multi-log increase in the number of colony-forming units in liver and spleen over time compared to WT animals. This increase is larger than what was reported under the same conditions for ISG15-deficient animals ^6^ and is reminiscent of *Listeria* infections in mice deficient in key immune signaling pathways necessary to control primary infection such as IFN-gamma ^80,81^. Further characterization of the immune response in the RNF213-deficient animals is both timely and relevant to assess whether a defect in cell autonomous immunity is underlying the increased susceptibility, as suggested by our *in vitro* data.

Co-localization of RNF213 with intracellular *Listeria* further supports a function of RNF213 as a restrictive host factor. Co-localization of proteins with intracellular bacteria is described as an antibacterial strategy known as xenophagy. Upon activation, specific E3 ubiquitin ligases, such as LRSAM1, SMURF1 or Parkin decorate invading pathogens with ubiquitin chains. Cytosolic adaptors proteins such as NBR1 or p62 recognize K48 or K63 ubiquitin chains and recruit LC3-II to capture the ubiquitinated bacteria into expanding autophagosomes ^82^. Since RNF213 has an active E3 ubiquitin ligase module, it is tempting to speculate that RNF213 conjugates ubiquitin chains to invading *Listeria* and facilitates its capture into autophagosomes. However, it remains to be determined if RNF213 is directly recruited to the surface of *Listeria* or rather to surrounding structures such as autophagic membranes on which the protein could oligomerize similar to what occurs on lipid droplets. Interestingly, wild-type bacteria evade xenophagy through cell-to-cell spread ^83^, thus RNF213 colocalization may be a related but distinct antimicrobial pathway that wild-type bacteria cannot evade. Alternatively, loss of bacterial and LD co-localization of the RNF213ΔC mutant could be analogous to what was shown for variants lacking of a functional ATPase core, i.e. that ubiquitination activity seems to be required for oligomerization and organelle targeting of RNF213 ^57^. Taken together, the data presented here provides the first evidence for co-localization of RNF213 with intracellular bacteria, classifying RNF213 as a new pathogen-directed E3 ubiquitin ligase.

Finally and most notably, the largest amount of information regarding RNF213 relates to MMD characterized by progressive stenosis of the internal carotid arteries and the secondary formation of a hazy network of basal collateral vessels in the brain ^84,85^. Allelic variations in the *RNF213* gene are the most important genetic risk factor to develop MMD, however, the functional role of RNF213 in MMD pathogenesis remains elusive ^86^. The best-characterized risk allele is the Asian-specific *RNF213* founder variant (p.R4810K) which dramatically increases the risk to develop MMD (e.g. >100-fold in Japan) and likely explains the higher MMD incidence in East-Asia. However, p.R4810K displays a strongly reduced penetrance since the incidence rate of MMD (e.g. 1/10,000 in the Japanese population) is much lower than the population p.R4810K carrier frequency (e.g. 1/50 Japanese carry the RNF213 R4810K risk allele) ^86^. This strongly suggests that additional genetic and/or environmental stimuli are required to induce MMD pathogenesis. Our study highlights that RNF213 plays a fundamental role as an antimicrobial host defense effector which could be inappropriately triggered during autoimmune responses, prior infection and/or inflammation in MMD patients ^87^. Indeed, previous work suggests that RNF213, like ISG15, is induced by IFN-I and pro-inflammatory cytokines ^88^. Here, we show that RNF213 oligomerization could depend on ISGylation of RNF213 and that the protein protects against infection with various pathogens. These data strongly support a role for infectious diseases as trigger to induce MMD in patients carrying RNF213 polymorphisms. It is conceivable that those patients have an impaired immune response to infection that could drive vascular fragility and increased vulnerability of vessels to hemodynamic stress and secondary insults. Together, the data presented here argues for a role of the immune response to infection in the development of MMD and should be fully explored in future studies of the etiology of this mysterious disease.

## ACKNOWLEDGEMENTS

We are grateful to Prof. Akio Koizumi for providing a 3xFLAG-RNF213 expression plasmid, to Prof. Pascale Cossart for sharing the *Listeria monocytogenes* EGD strain and *Listeria* EF-TU antisera, to Prof. Dr. Stacey Efstathiou for providing a GFP expressing HSV-1 strain and to Prof. Dr. Gary Cohen for providing the HSV-1 VP5 antisera. We thank Evelyn Plets and Annelies Van Hecke for help with the genotyping and *in vitro* infection experiments. A.B., K.P.K., and F.I. are supported by ERANET Infect-ERA BacVIRISG15. The German Research Foundation funded this project with grant KN590/7-1 to KPK, BE6335/6-1 to AB, CRC/TR167, project B16 to KPK and AB as well as CRC 1292, project 02 to AB. AB receives support from the Foundation for Experimental Biomedicine Zurich, Switzerland. CB is supported by International Max Planck Research School for Infectious Diseases and Immunology (IMPRS-IDI), Berlin. F.I. is supported by an Odysseus type 2 grant from the Research Foundation Flanders (G0F8616N). B.D. is supported by an Odysseus type 1 Grant of the Research Foundation Flanders (3G0H8318) and a starting grant from Ghent University Special Research Fund (01N10319). This work was supported by an NIH grant 1R35GM137961 to LR. XS acknowledges support from the FWO EOS project VIREOS and a BOF-UGent GOA project. N.C. is supported by the European Research Council (ERC) Consolidator Grant GlycoTarget (616966) and S.S. is supported by Research Grants Council, Hong Kong (17113019) and Wellcome Trust (220776/Z/20/Z).

---

**Supplementary Figure 1. RNF213 binds to premature, mature and non-conjugatable ISG15**

**(A-F)** HIV-1 GAG protein was genetically fused to wild-type ISG15, non-conjugatable ISG15AA or the precursor form of ISG15, and transiently expressed in HEK293T cells treated or not with IFN-I. Virus-like particles (VLPs) were purified, lysed and protein were digested into peptides for identification and quantification by LC-MS/MS. Volcano plots show the result of a t-test to compare VLPs containing ISG15 versus VLPs containing *E.coli* dihydrofolate reductase (eDHFR) as negative control (n=4 replicates). The fold change (in log2) of each protein is shown on the x-axis, while the statistical significance (−log P-value) is shown on the y-axis. Proteins outside the curved lines represent specific ISG15 interaction partners. Proteins identified as common ISG15 interaction partners in all six screens are indicated (n=29) and listed in Table S1. The volcano plot in (A) is identical to the volcano plot in Figure 1B.

**Supplementary Figure 2. Binding of RNF213 requires both N- and C-terminal domains of ISG15 (A-C)** HIV-1 GAG protein was genetically fused to ISG15AA (A), the ISG15 N-terminal (B) or the ISG15 C-terminal domain (C). Constructs were transiently expressed in HEK293T and Virotrap experiments were performed as in Fig. 1 and Supplementary Fig. 1 (n=3 replicates). Common ISG15 interaction partners from Supplementary Fig. 1 are annotated on the volcano plots and listed in Table S1. **(D)** Heatmap showing the expression level (log_2_ LFQ intensity) of ISG15, eDHFR, HIV-1 GAG and common ISG15 interaction partners from Supplementary Fig. 1 after non-supervised hierarchical clustering. On the right side, the same heatmap is shown with originally missing values colored in gray. All common ISG15 interactors including RNF213 bind to full length ISG15, but not to the C- or N-terminal domain alone.

**Supplementary Figure 3. ISG15 co-immunoprecipitation coupled to mass spectrometry**

**(A-C)** Immunoprecipitation using HA antibodies was performed from lysates of HEK293T or HeLa cells transfected with HA-ISG15AA or mock and treated with IFN-I for 24h or 48h. Pulled down proteins were digested with trypsin on the beads prior to their identification and quantification by LC-MS/MS. Volcano plots show the result of a t-test to compare ISG15 pull downs versus mock control (n=3 replicates). The fold change (in log2) of each protein is shown on the x-axis, while the statistical significance (−log P-value) is shown on the y-axis. Proteins outside the curved lines represent significantly enriched proteins. Except for ISG15 itself, no interacting proteins were identified in the ISG15 pull down experiments **(D)** Venn diagram showing the overlap between the common ISG15 interaction partners identified by Virotrap (n=29, Supplementary Fig. 1) and GST pull down in HEK293T cells (Figure 1D). Next to ISG15 itself, only RNF213 overlaps between those two type of experiments.

**Supplementary Figure 4. RNF213 binds non-covalently to ISGylated proteins but not to Ubiquitinated proteins**

**(A)** GFP immunoprecipitation was performed from lysates of HEK293T cells expressing eGFP-RNF213 or FLAG-eGFP in combination with HA-ISG15 and the ISGylation machinery (E1, E2, E3). A smear of ISGylated co-immunoprecipitated proteins was detected with eGFP-RNF213, but not with FLAG-eGFP.

**(B)** FLAG immunoprecipitation was performed in HEK293T cells expressing 3xFLAG-RNF213 or FLAG-eGFP in combination with HA-ISG15 and the ISGylation machinery (E1, E2, E3). After immunoprecipitation, beads were washed either with a buffer containing 1% Triton-X-100 or with a buffer containing 1% Triton-X-100 and 1 % SDS to remove ISGylated proteins that were non-covalently bound to FLAG-RNF213. **(C)** Workflow showing the strategy for co-immunoprecipitation of ISGylated proteins with separate expression of FLAG-RNF213 and ISGylated proteins. FLAG immunoprecipitation is performed from a lysate of HEK293T cells expressing 3xFLAG-RNF213. After binding of FLAG-RNF213, beads are washed and subsequently mixed with a lysate of HEK293T cells expressing HA-ISG15 and the ISGylation machinery (E1, E2, E3). In this way, we can exclude that (part of) the smear of co-immunoprecipitated ISGylated proteins is actually derived from ISGylated RNF213 itself. **(D)** FLAG immunoprecipitation was performed from lysates of HEK293T cells expressing 3xFLAG-RNF213, FLAG- RNF31 (HOIP) or FLAG-eGFP as control. According to the workflow in (C), after binding of the bait protein beads were mixed with a lysate of HEK293T cells expressing HA-ISG15 and the ISGylation machinery (E1, E2, E3). While FLAG-RNF213 was capable of pulling down ISGylated proteins, FLAG- RNF31 or FLAG-eGFP were not. **(E)** FLAG immunoprecipitation of 3xFLAG-RNF213, FLAG-RNF31 or FLAG-eGFP was performed as in (D), but after binding of the bait protein beads were mixed with a lysate of HEK293T cells expressing HA-Ubiquitin. FLAG-RNF31 efficiently pulled down ubiquitinated proteins, but FLAG-RNF213 and FLAG-eGFP did not.

**Supplementary Figure 5. RNF213 deficiency does not destabilize lipid droplets in macrophages.**

**(A)** THP-1 or primary human monocytes cells were cultured in the presence of 10 mM BSA-conjugated oleic acid. Lipid droplets (LDs)-enriched fractions were isolated by ultracentrifugation floatation assay on a sucrose step-gradient. Membrane bound fraction proteins and cytosolic proteins were isolated by ultracentrifugation sedimentation assay. SDS-PAGE and silver staining shows equal protein loading for each fraction (left panel). Immunoblotting reveals that RNF213 is mainly associated to lipid droplet. Immunoblotting against PLIN1, PLIN2, RAB18 and AUP1 shows efficient LD isolation. Immunoblotting against CNX and BIP was used as markers for the membrane fraction. Immunoblotting against GAPDH was used as a marker for the cytosolic fraction. **(B)** Representative images of BMDM cells derived from RNF213 +/+ or RNF213 -/- mice. Cells were treated with 10 ng/ml interferon-β, 200 µM mM BSA-conjugated oleic acid or both 24 h prior to fixation. Scale bars in the pictures are 10 microns. (C) At least 100 cells were used to count the volume (voxel, left panel) and number (right panel) of lipid droplets per cell (right panel) from five different fields containing each 20 to 30 cells. Quantification was performed with the Volocity software. No differences in volume or number of lipid droplets were observed between RNF213 +/+ and RNF213 -/- cells (AVG ± SEM, n=5 fields, two-tailed unpaired t-test), indicating that RNF213 deficiency does not lead to reduced stability of lipid droplets in macrophages.

**Supplementary Figure 6. IFN-I induction and regulation of RNF213 and ISG15 in human cell lines**

(**A**) HeLa, A549 or THP-1 cells were either untreated or treated with the indicated type of interferon (α/β/γ) for 48h prior to lysis and immunoblotting against RNF213, ISG15 and tubulin as loading control. RNF213 and ISG15 are co-induced, primarily by IFN-α and β. (**B**) HeLa cells were treated or not with interferon-β prior to lysis and distribution of RNF213 between the soluble and membrane-associated fraction was quantified by immunoblotting. Purity of the soluble and membrane associated fractions was evaluated using GAPDH and Ribophorin I as respective marker proteins (AVG ± STDEV, n=4 replicates). Interferon treatment led to a non-significant decrease and increase of RNF213 in the soluble and membrane fractions, respectively. (**C**) Immunoblots against UBE1L and tubulin as loading control confirmed knockdown of UBE1L in the experiment shown in Fig. 3B.

**Supplementary Figure 7. RNF213 counteracts in vitro infection**

**(A)** A549 cells were infected with RSV-A2 for up to six days at MOI 0.001. Immunoblotting against RSV- G confirmed infection, while pSTAT1 indicated type-I interferon signaling especially 2 to 4 days post infection. Tubulin was used as loading control. **(B)** HeLa cells were infected with coxsackievirus B3 (CVB3) up to 48 h at MOI 0.01. Immunoblotting against coxsackie viral protein VP1 confirmed infection, while the absence of pSTAT1 indicated no induction of type-I interferon signaling. **(C)** HeLa cells were infected with *Listeria monocytogenes* EGD up to 48 h at a multiplicity of infection (MOI) of 25. Immunoblotting against *Listeria* EF-TU confirmed infection, while phosphorylated STAT1 (pSTAT1, Y701) showed a slight induction of type-I interferon signaling 24 h post infection. Tubulin was used as loading control. **(D)** HeLa cells were infected with *Listeria* for 16 h at MOI 25. The percentage of bacteria inside HeLa cells transfected with a pool of siRNAs against RNF213 (siRNF213 pool, used in all other experiments) or individual siRNAs against RNF213 (siRNF213 D, siRNF213 INV6645, siRNF213 INV4009) is shown relative to siSramble-transfected cells (AVG ± SEM, n=3 independent experiments, two-tailed unpaired t-test, * p < 0.05, ** p < 0.01 and *** p < 0.001). **(E)** HeLa cells were infected with *Listeria* for 16 h at MOI 25. 48 h prior to infection, cells were transfected with a pool of siRNAs targeting RNF213 (siRNF213) or a pool of non-targeting scrambled siRNAs (siScramble) as control. Additionally, 24 h prior to infection cells were transfected with a plasmid encoding HA-ISG15 or with an empty vector (mock) as control. Intracellular *Listeria* were quantified after serial dilution by counting colony-forming units (CFUs) in a gentamycin assay. The percentage of intracellular bacteria relative to mock transfected cells is shown (AVG ± SEM, n=3 independent experiments, two-tailed unpaired t-test, ** p < 0.01, *** p < 0.001).

**Supplementary Figure 8. RNF213 knockout mice are strongly susceptible to Listeria infection**

**(A)** Generation of C57BL/6J RNF213 KO mice. The largest exon (exon 28) of the Rnf213 gene is 3603 bp and was targeted by 2 gRNAs, Rnf213_gRNA1 with protospacer sequence 5’ CAGAGCTTCGGAACTTTGCT 3’ and Rnf213_gRNA2, with protospacer sequence 5’ TGTGCCCCTCATCAACCGTC 3’. The distance between the Cas9 cut sites of both gRNAs is 2842 bp. Screening for the deletion between both gRNAs was done with primers Rnf213_FW1 (5’ AGTTTCTTGATCTCTTCCCC 3’) and Rnf213_Rev6 (5’ CTCCTCCGTCAGATCCCTA 3’) generating a wild type PCR fragment of 3363 bp and a fragment of 523 bp in case of an exact deletion between both Cas9 cut sites. Several founders showed a deletion band. Founder 290-7 was further analyzed and showed a deletion of 2854 bp, resulting in a frameshift and premature stopcodons resulting in mouse line B6J- RNF213^em1Irc^. This line was used for further breeding and the generation of RNF213 -/- and RNF213 +/+ littermate control mice. **(B-E)** RNF213 -/- and RNF213 +/+ littermates were infected intravenously with 5 × 10^5^ *Listeria monocytogenes* EGD. Liver and spleen were isolated following 72 h of infection, and CFUs per organ were counted by serial dilution and replating; dots and squares depict individual animals (AVG ± SEM, Mann–Whitney test, ** p < 0.01 and *** p < 0.001). **(B-C)** Six mice per genotypes were used in experiment 2. **(D-E)** Seven RNF213 -/- and eight RNF213 +/+ mice were used in experiment 3 (note that the results of experiment 1 are shown in Fig. 7). Thus, in three independent experiments, 72 h post infection RNF213 -/- animals showed significantly higher bacterial loads in liver (∼1 log) and spleen (∼5 logs) compared to WT littermate controls.

**Supplementary figure 9. The RNF213 E3 module is required for lipid droplet localization**

(A) Representative images of HeLa cells transfected with eGFP-RNF213 or eGFP-RNF213ΔC for 72 hours. Following transfection cells were left untreated for 18 h and then fixed. Scale bars in the pictures and insets are respectively 10 microns and 0.5 microns. While eGFP-RNF213 adopts a spherical pattern reminiscent of lipid droplet localization reported by Sugihara et al., 2019, eGFP-RNF213ΔC shows a diffuse cellular staining. (B) Representative images of an independent experiment in which HeLa cells were transfected with eGFP-RNF213ΔC and counterstained for lipid droplets. 72 hours post transfection cells were left untreated for 18h and then fixed. Scale bars in the pictures are 10 microns. Again, eGFP- RNF213ΔC was spread throughout the cell without co-localization to lipid droplets.

## METHODS

### Antibodies

The following primary antibodies were used for immunoblotting: mouse monoclonal anti-ISG15 (F-9, sc- 166755, Santa Cruz Biotechnology), rabbit monoclonal anti-ISG15 (Abcam [EPR3446] ab133346), mouse monoclonal anti-ubiquitin (P4D1, #sc-8017, Santa Cruz Biotechnology), mouse monoclonal anti-α-tubulin (B-7, #sc-5286, Santa Cruz Biotechnology), mouse monoclonal anti-GST (B-14, #sc-138, Santa Cruz Biotechnology), rabbit polyclonal anti-GAPDH (FL-335, #sc-25778, Santa Cruz Biotechnology), rabbit polyclonal anti-HA-tag (#H6908, Merck), rabbit polyclonal anti-RNF213 (#HPA003347, Merck), mouse monoclonal anti-RNF213 (clone 5C12; Santa Cruz Biotechnology), mouse monoclonal anti-FLAG-tag (M2, #F3165, Merck), rabbit polyclonal anti-LC3B (#PA146286, ThermoFisher scientific), rabbit polyclonal anti-tubulin-α (#ab18251, Abcam), anti-p24 GAG (#ab9071, Abcam), rabbit polyclonal anti- PLIN1 (#ab3526, Abcam), rabbit anti-PLIN2 (#ab108323, Abcam), rabbit polyclonal anti-Rab18 (#ab119900, Abcam), anti-AUP1 (Zhang et al., 2018), rabbit polyclonal anti-BiP (Abcam ab21685), rabbit polyclonal anti-Calnexin (Abcam ab10286), rabbit polyclonal anti-ATGL (Cell Signaling Technology #2138). *Listeria* EF-Tu was immunoblotted with rabbit polyclonal antisera raised against α-EF-Tu (Archambaud et al., 2005). Herpes Simplex virus-1 VP5 was immunoblotted with rabbit polyclonal anti-NC-1 antiserum specific for HSV-1 VP5 (Cohen et al., 1980). Respiratory syncytial virus RSV-G was immunoblotted with commercially available goat polyclonal anti-RSV serum (#AB1128, Merck). Coxsackievirus VP1 was immunoblotted with commercially available mouse monoclonal anti-VP1 (3A8, Mediagnost). Aforementioned primary antibodies were revealed using goat polyclonal anti-mouse-IgG (IRDye® 800CW, Li-COR), goat polyclonal anti-rabbit-IgG (IRDye® 800CW, Li-COR), goat polyclonal anti-mouse-IgG (IRDye® 680RD, Li-COR) or goat polyclonal anti-rabbit-IgG (IRDye® 680RD, Li-COR), except for anti-RSV serum which was revealed with secondary anti-goat (#sc-2020, Santa Cruz biotechnology). For microscopy, mouse monoclonal anti-ISG15 (3E5, #sc-69701, Santa Cruz biotechnology) was used and revealed with goat polyclonal anti-mouse IgG secondary Superclonal™ Recombinant Secondary Antibody, Alexa Fluor 647 (#A28181, Thermo Fisher Scientific).

### Cell culture

Hek293T cells originate from (Lin et al., 2014) while HeLa cells (ATCC® CCL-2™), A549 cells (ATCC® CCL-185™) THP-1 cells (ATCC® TIB-202™) and primary human CD14+ monocytes (ATCC® PCS- 800-010^™^) were purchased from ATCC. HEK293T and A549 cells were cultured in DMEM medium (#31966047, Thermo Fisher Scientific) supplemented with 10% fetal bovine serum (FBS, #10270106, Thermo Fisher Scientific). HeLa cells were maintained in MEM medium (#M2279, Merck) supplemented with 10% FBS, 1% glutamax (#35050038, Thermo Fisher Scientific), 1% non-essential amino acids (#11140035, Thermo Fisher Scientific), 1% sodium pyruvate (#11360039, Thermo Fisher Scientific) and 1% hepes (#15630056, Thermo Fisher Scientific). THP-1 cells were maintained in RPMI 1640 medium (#61870044, Thermo Fisher Scientific) supplemented with 10% FBS and 2 mM L-glutamine (#25030024, Thermo Fisher Scientific). THP-1 cells were differentiated to macrophages by complementing the media with 10 ng/ml phorbol 12-myristate 13-acetate (#P8139, Merck) for seven days, followed by rest for 24 h in medium RPMI 1640 medium supplemented with 10% FBS, 2 mM L-glutamine. Primary human CD14+ monocytes were maintained in Hank’s Balanced Salt Solution (HBSS, #14175095, Thermo Fisher Scientific) supplemented with 10% FBS. HEK293T, HeLa and A549 cells were authenticated by Eurofins.

### Plasmid transfection

Plasmid transfection was performed with Polyethylenimine (PEI, #23966-1, Polysciences) as transfection reagent at a ratio PEI/cDNA of 5:1 (w/w). Plasmids were used at a final concentration of 1 μg DNA/10^6^ Hek293T cells or 1 μg DNA/5×10^5^ HeLa cells. For microscopy, HeLa cells were reverse-transfected at the day of seeding with 17µg of plasmid either coding for eGFP-RNF213 or eGFP-RNF213-ΔC using 51 µL of Fugene® HD (#E2311, Promega) for a ratio of 1:3 (µg cDNA / µL transfection reagent). HeLa cells were cultured for 48 h and moved to a 6 wells-plate with coverslip for microscopy. The following plasmid were used: pMD2.G (VSV-G), pcDNA3-FLAG-VSV-G, pMET7-GAG-EGFP, pMET7-GAG-ISG15GG, pMET7-GAG-ISG15AA, pMET7-GAG-ISG15-precursor, pGL4.14-3xFLAG-RNF213, pGL4.14- 3xFLAG-RNF213ΔC, pGL4.14-3xFLAG-RNF213_R4810K_, pMet7-FLAG-C-domain-RNF213, pRK5-HA- RNF4, pCMV3-FLAG-HOIP (RNF31), pMET7-Flag-eGFP, pcDNA3.1-HA-ISG15GG, pccDNA3.1-HA-ISG15AA, pSVsport (Mock plasmid), pcDNA3.1-hUbe1L (E1), pcDNA3.1-Ubch8 (E2) and pTriEx2- hHERC5 (E3), pDEST-eGFP-RNF213 and pDEST-eGFP-RNF213ΔC. The cDNA sequences of all aforementioned plasmids are available on addgene.org or listed in Table S2.

### SiRNA transfection

A commercially available siRNA pool was used to knockdown human RNF213 (#M-023324-02, GE Healthcare Dharmacon) in figure 2, 3, 4 and 5. As control siRNA treatment, a non-targeting scramble siRNA (#D-001210-01-05, GE Healthcare Dharmacon) was used in the experiments shown in figure 3 and supplementary figure 6C while a pool of four scrambled siRNAs (#D-001206-13-05, GE Healthcare Dharmacon) was used in the experiments shown in figure 2, 4 and 5. In all experiments, siRNAs were transfected with DharmaFECT transfection reagent (#T-2001-02, GE Healthcare Dharmacon) according to the instructions of the manufacturer. For supplementary fig. 5A, the siRNA D-023324-05 (GE Healthcare Dharmacon) and Stealth RNAis (#HSS126645, #HSS184009, Thermo Fisher Scientific) were also transfected in HeLa cells to knockdown RNF213 with DharmaFECT and Lipofectamine 3000 transfection reagent (#L3000008, Thermo Fisher scientific), respectively. For figure 4, a commercially available siRNA pool was used to knockdown ISG15 (#M-004235-04-005, GE Healthcare Dharmacon). In these experiments, a reverse siRNA transfection protocol was adopted to knockdown the expression of ISG15 and RNF213 genes in HeLa cells prior to *Listeria* infection. A single siRNA, custom produced by Dharmacon, was used to knockdown RSV-N expression as described in (Alvarez et al., 2009). Immunoblotting assays were conducted to confirm reduction of protein expression levels.

### Generation of knockout cell lines

The RNF213 knockout HeLa cell line was generated by using a CRISPR/Cas9 approach. Target sequences were selected by CRISPOR (CRISPOR.org, (Concordet and Haeussler, 2018). Oligonucleotides (5’CACCGGAGGCAGCCTCTCTCCGCAC and 5’AAACGTGCGGAGAGAGGCTGCCTCC; 5’CACCGTGCAGCCCCCATAGCAGGTG and 5’AAACCACCTGCTATGGGGGCTGCAC) were synthesized by ID&T (Leuven, Belgium) and cloned into the pSpCas9(BB)-2A-Puro plasmid (pXP459, Addgene #48139). HeLa cells were co-transfected with the two RNF213-targeting plasmids with Lipofectamine 3000 (L3000008, Thermo Fisher scientific) as described above. Cells were selected with 2 µg/mL with Puromycin (#P8833, Merck) for 48 h. Cells were diluted and plated in 96-well plate for single clone selection. The absence of RNF213 protein expression was confirmed by immunoblotting. The ISG15 knockout HeLa cell line was generated as previously described (Kespohl et al., 2020).

### SDS-PAGE and immunoblotting

Cells were lysed in 2x Laëmmli buffer containing 125 mM Tris-HCl pH 6.8, 4% SDS, 20% glycerol, 0,004% Bromophenol blue supplemented with 20 mM DTT. Protein samples were boiled for 5 minutes at 95°C and sonicated prior to SDS-PAGE. Samples were loaded on 4-20% polyacrylamide gradient gels (#M42015, Genescript), 4–15% Mini-PROTEAN TGX Gels (#4561084, Biorad), 3-8% Criterion XT tris-acetate gel (#3450130, Biorad) or 4-15% Criterion TGX gel (#5671083, Biorad) according to the guidelines of the manufacturer. For detection of RNF213, proteins were separated on a 3-8% Criterion XT tris-acetate gel (Biorad) or 4-15% Criterion TGX gel according to the instructions of the manufacturer. Proteins were transferred to PVDF membrane (#IPFL00010, Merck) for 3 hours at 60 V with Tris/Boric buffer at 50 mM/50 mM. Membranes were blocked for 1 hour at room temperature (RT) with blocking buffer (#927- 50000, LI-COR) and incubated with primary antibodies overnight at 4 °C diluted to 1:1000 in TBS. The next day, membranes were washed three times for 15 minutes with TBS-Tween 0.1% (v/v) buffer and further incubated at RT for 1 h with the appropriate secondary antibody. Membranes were washed twice with TBS-tween 0.1% (TBS-T) and once with TBS prior to detection. Immunoreactive bands were visualized on a LI-COR-Odyssey infrared scanner (Li-COR).

### Lipid droplets isolation and cellular fractionation

THP-1 cells treated or not with IFN-I (100 U/mL; 8-10 h) were washed and scraped in ice-cold hypotonic disruption buffer containing 20 mM tricine and 250 mM sucrose pH 7.8, 0.2 mM PMSF followed by homogenization in a chilled glass homogenizer. THP-1 lysates underwent a nitrogen bomb at a pressure of 35 bar for 15 min on ice to complete cellular disruption. Lysates were centrifuged at 3,000 x g for 10 min at 4°C and the post-nuclear supernatants was loaded at the bottom of 13-mL centrifuge tubes and overlaid with ice-cold wash buffer containing 20 mM HEPES, 100 mM KCl and 2 mM MgCl2 at pH 7.4. Tubes were centrifuged in a swing-out rotor at 182,000 x g at 4°C for 1 h. Lipid droplets were collected from the top of the gradient. Lipid droplet-enriched samples were centrifuged at 20,000 x g for 5 min at 4°C to separate the floating lipid droplets from the aqueous fraction. Lipid droplets were washed three times by adding 200 µL ice-cold wash buffer followed by centrifugation at 20,000 x g for 5 min at 4°C. Lipid droplets were resuspended in 100 µL 2x Laëmmli buffer and further processed for immunoblot analysis as described above. To isolate the membrane fraction, THP-1 lysates were centrifuged at 3,000 x g for 10 min at 4°C and the pellet was washed three times by resuspension with 1 mL of ice-cold wash buffer followed by centrifugation. The insoluble fractions were resuspended in 2x Laëmmli buffer and further processed for immunoblot analysis as described above. To isolate the cytosolic fraction, THP-1 lysates were centrifuged at 3,000 xg for 10 minutes and 1 mL aliquot was sampled from the middle of the lysate. The 1mL-aliquot was further centrifuged at 270,000g for 1 h at 4 °C in a microcentrifuge tube using a TLA100.3 rotor. The supernatants were collected and mixed to a final 2x Laëmmli buffer prior further processing for immunoblot analysis as described above. For cellular fractionation in HeLa cells, cells were grown on 100 cm^2^ petri dishes and either untreated or treated for 24 h with interferon-β at 10 ng/ml (#11343524, Immunotools). Cells were processed as described in (Weber et al., 2017). Briefly, cells were detached with trypsin, washed with PBS and lysed in a buffer containing 300 mM sucrose, 5 mM Tris-HCl, 0.1 mM EDTA, pH 7.4 and a protease inhibitor cocktail (Roche, cOmplete, Mini, EDTA-free tablet) followed by homogenization in a chilled glass homogenizer. Lysates were centrifuged at 800 x g for 8 min at 4°C followed by separation of the soluble and membrane fractions by a second centrifugation step at 20,000 x g for 120 min at 4°C. The membrane-associated fraction was re-dissolved in lysis buffer to a volume equal to the volume of the soluble fraction. Samples were mixed with Laëmmli buffer and processed for immunoblotting as described above.

### Glycerol gradient analyses

Lysates prepared as described for the purification of lipid droplets were loaded onto a glycerol gradient (10%–40% (w/v). The gradients were centrifuged at 237,000 x g for 20 h at 4°C, using a SW 55 Ti rotor and Beckman L-80 ultracentrifuge. Twenty fractions for each sample were collected in 2 mm increments using a BioComp Piston Gradient fractionator. Fractions were concentrated by TCA precipitation and analyzed by immunoblotting as described above. For immunoprecipitations, each gradient fraction was first desalted over Amicon filter to remove glycerol and resuspended in 1 ml of resuspension buffer containing 20 mM HEPES, 100 mM KCl and 2 mM MgCl2 at pH 7.4. Resuspended fractions were pre-cleared with Protein G-agarose beads by incubation for 1 h at 4°C. Immunoprecipitation of Rnf213 was performed from pre-cleared fractions by incubation with RNF213 antibody for 3 hours at 4°C with end-over-end rotation. Beads were washed twice in 1 mL of resuspension buffer by centrifugation and samples were eluted into 1X Laëmmli buffer by heating at 60°C and processed for immunoblotting as described above.

### In vitro infection with Listeria monocytogenes

*Listeria monocytogenes* (EGD BUG600 strain) was grown in brain heart infusion (BHI) medium at 37°C. *Listeria* were cultured overnight and then sub cultured 1:10 in BHI medium for 2 h at 37°C. Bacteria were washed three times in PBS and resuspended in medium without FBS prior to infection. HeLa cells were grown in 6-well plates and infected with *Listeria* at a multiplicity of infection (MOI) of 25. Right after infection, plates were centrifuged at 1,000 x g for 1 min followed by incubation for 1 hour at 37°C to allow entry of the bacteria. Afterwards, cells were washed two times with PBS and then grown in MEM medium with 10% FBS, 1% glutamax, 1% non-essential amino acids, 1% sodium pyruvate, 1% hepes, supplemented with 40 µg/mL of gentamicin to kill extracellular bacteria. For immunoblotting, infected cells were washed with ice-cold PBS, lysed in 2x Laëmmli buffer and further processed as described above. To count the number of intracellular bacteria, HeLa cells were washed and lysed with miliQ water to release intracellular bacteria. Colony Forming Units (CFUs) were determined by serial dilution and plating on BHI agar.

### In vivo infection with Listeria monocytogenes

*Listeria monocytogenes* (EGD BUG600 strain) was grown as described above. Female and Male C57BL/6 mice (RNF213+/+ or RNF213−/−) between 8 and 12 weeks of age were infected intravenously by tail vein injection with 5 × 10^5^ bacteria per animal. Mice were sacrificed 72 h following infection. CFUs per organ (liver or spleen) were enumerated by serial dilutions after tissue dissociation in sterile saline. The animals were housed in a temperature-controlled environment with 12 h light/dark cycles; food and water were provided ad libitum. The animal facility operates under the Flemish Government License Number LA1400536. All experiments were done under conditions specified by law and authorized by the Institutional Ethical Committee on Experimental Animals (Ethical application EC2019-080).

### *In vitro* infection with Herpes Simplex Virus Type 1 strain C12 (HSV-1 C12)

HSV-1 C12 is a recombinant HSV-1 strain SC16 containing a cytomegalovirus (HCMV) IE1 enhancer/promoter-driven enhanced green fluorescent protein (EGFP) expression cassette in the US5 locus (kindly provided by Stacey Efstathiou (University of Cambridge, Cambridge, UK). HSV-1 C12 was propagated in Vero cells and titrated by plaque assay with Vero cells according to L. Grosche et al., 2019. HeLa cells were grown in 96-well plates and infected with HSV-1 C12 at MOI 0.1 (figure 4A-C) or MOI 0.05 (figure 4D-E). Cells were treated either without or with 1000 U/mL IFNα2a (**#** 11343504, Immunotools). Cells were maintained in 2% FBS FluoroBrite DMEM Media. GFP fluorescence was monitored at 24 h intervals using the Infinite® 200 PRO Reader and Tecan iControl Software (Tecan Life Sciences). Each individual measurement (n=4) was normalized to the mean value of the uninfected wells (n=4). The relative area under the curve from 0 to 72 h post infection was calculated for each growth curve according to (Crameri et al., 2018). For immunoblot analysis, cells 24 h p.i were washed with ice-cold PBS, lysed in 2x Laëmmli buffer and further processed as described above.

### *In vitro* infection with coxsackievirus B3 (CVB3)

CVB3 Nancy strain was propagated in Green Monkey Kidney Cells and quantified by plaque assay in HeLa cells as described below. HeLa cells were grown in 6-well plates and infected with CVB3 (MOI 0.01) in serum free medium. After 1 h, the CVB3 inoculum was replaced and HeLa cells were cultured in MEM medium with 10% FBS, 1% glutamax, 1% non-essential amino acids, 1% sodium pyruvate and 1% hepes. For immunoblotting, the cells were washed with ice-cold PBS, lysed in 2x Laëmmli buffer and further processed as described above. For relative quantification upon knockdown of RNF213, intensity of the VP1 band was normalized to its intensity in the siScramble condition. For qRT-PCR, cells were lysed in 500 µl TRIzol™ Reagent. RNA was extracted from the aqueous phase after chloroform addition and precipitated by addition of isopropyl alcohol. 250 ng RNA per sample was used for reverse transcription to cDNA. For relative quantification, the ΔΔCt method was used with Hypoxanthine Phosphoribosyl transferase (HPRT) as housekeeping gene. Plaque forming units (PFU)/mL were determined by serial dilution on confluent HeLa cells.

### *In vitro* infection with respiratory syncytial virus (RSV)

RSV-A2, an A subtype of RSV (ATCC, VR-1540, Rockville), was propagated in HEp-2 cells and quantified by plaque assay with A549 cells as described below. A549 cells were grown in 6-well, 48-well or 96-well plates respectively for immunoblot, titration or plaque assay. A549 cells were infected in serum-free medium with RSV-A2 at MOI 0.001, MOI 0.02 or MOI 0.005 for the respective experiment. After 4 h, the RSV inoculum was replaced and A549 cells were cultured in DMEM medium with 10% FBS. For immunoblot analysis, cells were washed with ice-cold PBS, lysed in 2x Laëmmli buffer and further processed as described above. In the plaque and titration assay, A549 cells were treated either without or with 10 ng/mL interferon-β (#11343524, Immunotools) for 48 h. For plaque assays, the DMEM medium with 10% FBS was supplemented with 0,6% (w:v) avicel RC-851 (FMC biopolymers) and the cells were fixed 6 days post RSV infection with 4% paraformaldehyde (PFA) in PBS and permeabilized with 0,2% Triton-X100 in PBS. RSV plaques were stained with a polyclonal goat anti-RSV serum and secondary anti-goat IgG. Plaques were visualized by TrueBlue^TM^ Peroxidase substrate (5510, Sera Care). Each plaque was quantified in Fiji. In titration assays, Plaque Forming Units (PFU)/mL were determined by serial dilution on confluent A549 cells. Supernatants were collected and mixed (1:1) with a 40% sucrose solution in HBSS, snap frozen and stored at -80°C.

### BMDM’s

Female and Male C57BL/6 mice (RNF213+/+ or RNF213−/−) between 8 and 12 weeks of age were used to isolate bone marrow cells from femurs. Isolated bone marrow cells were cultured for 7 days at a density of 105 cells/mL in DMEM/F-12 medium, supplemented with 10% FBS, 10 units/ml penicillin, 10 μg/mL streptomycin, and 20 ng/mL murine M-CSF at 37°C in a humidified atmosphere with 5% CO_2_. On day 3, fresh medium containing 40 ng/ml M-CSF was added. Cells were further differentiated for 4 days in M-CSF containing medium.

### Microscopy and image processing

HeLa cells were plated on coverslips the day prior to an experiment. Cells were either not treated or infected as described above with *Listeria monocytogenes* strain EGD or infected with the same strain stably expressing mCherry for either 24 hours at MOI 25 or for 18 h at MOI 5, respectively. Cells were washed with PBS and subsequently fixed with 4% PFA (Electron Microscopy Sciences, Hatfield, Pennsylvania) in PBS for 20 minutes at room temperature, then permeabilized with 0.5% Triton in PBS. Only for the figure 6A and supplementary figure 9B, cells were then counterstained for lipid droplets (HCS LipidTOX™ Red Neutral Lipid Stain, Thermo Fisher Scientific). Coverslips were mounted using DAPI Fluoromount-G® (SouthernBiotech, USA), and images were acquired using an inverted wide-field fluorescence microscope (Axio Observer 7, Carl Zeiss Microscopy, Germany) equipped with an Axiocam 506 mono camera and the software ZEN 2.3 Pro. Images were deconvoluted using Zen 3.1 (Blue Edition) for optimal resolution. For figure 6B, the analysis of the number of lipid droplet per cell and the colocalization of eGFP-RNF213 with Listeria was performed using Fiji (ImageJ) software. For supplementary figure 9B-C, BMDMs used for microscopy were seeded in IBIDI μ-Slide 8 Well chambers and treated with oleic acid conjugated to BSA (molar ratio 5:1 in PBS) at a concentration of 200µM. Interferon-β was used at 10 ng/mL. BMDMs were fixed in PBS with 4% PFA for 20 min at RT. LHCS LipidTOX™ Red Neutral Lipid Stain, Thermo Fisher Scientific) and Hoechst were used to visualize lipid droplets and nuclei respectively. Cells were mounted using Prolong Diamont Mounting Medium (Thermo, USA). Images were acquired using a confocal laser scanning microscope (LSM880 with Airyscan, Carl Zeiss Microscopy, Germany) and the software ZEN 2.3 Pro. And deconvoluted using Zen 3.1 (Blue Edition) for optimal resolution. The number and relative area of LDs were analyzed using Volocity software.

### Virotrap sample preparation for LC-MS/MS and immunoblotting analysis

Virotrap experiments were performed as described in (Titeca et al., 2017). Briefly, the day before transfection 10^6^ Hek293T cells (authenticated) were seeded in four T75 flasks for each condition. Cells were then transfected with four different bait proteins, each fused via their N-terminus to HIV-1 GAG: mature ISG15 (ending on -LRLRGG), mature non-conjugatable ISG15 (ending on -LRLRAA), full length ISG15 precursor, and eDHFR (dihydrofolate reductase from *Escherichia coli*) as control. VSV-G and FLAG-VSV-G were co-transfected to allow single-step purification of the produced particles. The day after transfection, these cells were treated with interferon-α for 24 h (10 ng/mL; #11343596, Immunotools) during the particle production phase. Supernatant was harvested 40 h after transfection followed by centrifugation and filtration (0.45 µM; SLHV033RS, Merck Millipore) to remove cellular debris. Virotrap particles containing protein complexes were purified in a single step using biotinylated anti-FLAG BioM2 antibody (#F9291, Merck) and Dynabeads MyOne Streptavidin T1 Beads (#65601, ThermoFisher scientific), and were consecutively eluted by competition using FLAG peptide (#F3290, Merck). For immunoblot analysis, samples containing the purified VLPs were mixed 1:1 with 4X Laëmmli buffer buffer complemented with 20mM DTT further analyzed as described above. After FLAG elution, samples were processed with Amphipols A8-35 (#A835, Anatrace), digested using trypsin (V5111, Promega), and acidified. Each condition (4 baits, +/- interferon-α) was analyzed in quadruplicate, leading to a total of 32 samples for LC-MS/MS analysis. Peptides were purified on Omix C18 tips (Agilent), dried and re-dissolved in 20 µl loading solvent A (0.1% trifluoroacetic acid in water/acetonitrile (ACN) (98:2, v/v)) of which 2.5 µl was injected for LC-MS/MS analysis.

### GST pull down for LC-MS/MS and immunoblotting

HEK293T, HeLa or differentiated THP-1 cells were grown at ∼80-90% confluence in 15 cm culture dishes (1 petri dish/sample). HEK293T cells were treated with interferon-α while HeLa and THP-1 cells were treated with interferon-β, at 10 ng/mL for 24 h (#11343596, #11343524, Immunotools). Each condition (2 baits, 3 cell lines) was analyzed in triplicate, leading to a total of 18 samples for LC-MS/MS analysis. Cells were washed three times with PBS with Ca^2+^ and Mg^2+^ and scraped in 1 mL lysis buffer containing 50 mM Tris.HCl (pH 8.0), 150mM NaCl, 1% triton-x-100 (v/v), 1 mM PMSF and a protease inhibitor cocktail (Roche, cOmplete, Mini, EDTA-free tablet, 4693159001). Samples were incubated 60 minutes on ice or 20 minutes under end-over-end agitation at 4°C. Lysates were cleared by centrifugation for 15 minutes at 16,000 x g at 4°C to remove insoluble components. 15 µL magnetic glutathione beads (Pierce ThermoFisher scientific) were incubated with 4 µg of purified GST-tagged ISG15 (R&D systems, UL-600-500) or GSTP1 (#G5663, Merck) under agitation. For immunoblotting, glutathione beads were also decorated with GST-tagged-SUMO1 (R&D systems, UL-710-500) or GST-tagged-Ubiquitin (R&D systems, U-540-01M). Incubation was performed overnight at 4°C in 500 µL 50 mM Tris.HCl, 150 mM NaCl, 1% triton-x-100 (v/v), 1 mM DTT after which decorated glutathione beads were blocked by addition of BSA to a final concentration of 2% (w/v) and incubation for an additional hour at 4°C. Then, the cleared lysates were incubated overnight at 4°C with the decorated glutathione beads under agitation to allow binding of proteins to ISG15. The following day, the beads were precipitated with a magnetic stand, washed once with 1mL wash buffer containing 50 mM Tris.HCl (pH 8.0), 150mM NaCl, 1% triton-x-100 (v/v). For immunoblotting, the beads were washed two extra times with 1mL wash buffer for a total of three washes. Beads were mixed with 60µL 2X Laëmmli buffer complemented with 20mM DTT and boiled at 95°C for 10 minutes. Beads were precipitated with a magnet and the supernatants were further analyzed by immunoblotting as described above. In the LC-MS/MS analysis, beads were washed three extra times with 1 mL trypsin digestion buffer containing 20 mM Tris HCl pH 8.0, 2 mM CaCl2. Washed beads were re- suspended in 150 µl digestion buffer and incubated for 4 hours with 1 µg trypsin (Promega) at 37 °C. Beads were removed, another 1 µg of trypsin was added and proteins were further digested overnight at 37 °C. Peptides were purified on Omix C18 tips (Agilent), dried and re-dissolved in 20 µl loading solvent A (0.1% trifluoroacetic acid in water/acetonitrile (ACN) (98:2, v/v)) injected for LC-MS/MS analysis. In the experiment with HeLa cells, 15 µL of samples was injected while 5µL was injected for the experiment with THP-1 and Hek293T cells.

### Immunoprecipitation (IP) for LC-MS/MS and immunoblotting

HEK293T and HeLa cells were grown at ∼80-90% confluence in 15 cm culture dishes (1 petri dish/sample). In the IP-immunoblotting experiments, cells were transfected either with FLAG-tagged-eGFP or FLAG- RNF213 plasmids, plus, the ISG15 conjugation machinery with HA-tagged-ISG15 matured, UBA7, UBCH8 and HERC5 plasmids. In the IP-MS experiment, cells were transfected either with plasmid coding for HA-tagged-ISG15AA or a mock plasmid. HEK293T cells were treated with interferon-α at 24h or 48h while HeLa cells were treated with interferon-β at 24h. Intereferon-α/β treatment were performed at 10 ng/mL (#11343596, #11343524, Immunotools). Each condition (2 baits, 3 types of interferon treated cells) was analyzed in triplicate, leading to a total of 18 samples for LC-MS/MS analysis. Cells were washed three times with PBS with Ca^2+^ and Mg^2+^ and scraped in 1.5 mL lysis buffer containing 50 mM Tris.HCl (pH 8.0), 150mM NaCl, 1% triton-x-100 (v/v), 1 mM PMSF and a protease inhibitor cocktail (Roche, cOmplete, Mini, EDTA-free tablet, 4693159001). Samples were incubated 60 minutes on ice or 20 minutes under end-over-end agitation at 4°C. Lysates were cleared by centrifugation for 15 minutes at 16,000 x g at 4°C to remove insoluble components. Supernatants were incubated with Pierce anti-HA magnetic beads (#88836, Pierce ThermoFisher scientific) or anti-FLAG magnetic beads (#M8823, Merck). Lysates were incubated at 4°C under agitation overnight. The beads were precipitated with a magnetic stand and subsequently washed once with wash buffer containing 50mM Tris.HCl (pH 8.0), 150mM NaCl, 1% triton-x-100 (v/v). In the IP-immunoblotting experiments, the beads were washed two extra times with wash buffer. Beads were mixed with 60µL 2X Laëmmli buffer and heated at 60°C for 10 minutes. Beads were precipitated with a magnet and the supernatants were further analyzed by immunoblot analysis as described above. In the IP-MS experiment, the beads were washed three extra times with trypsin buffer containing: 20 mM Tris HCl pH 8.0, 2 mM CaCl_2_. The proteins were on-bead digested with 1 µg trypsin (Promega) for 4 hours at 37 °C under agitation. Then, the beads were precipitated and the supernatants were mixed with 1µg of trypsin and further digested overnight at 37°C under agitation. Peptides were desalted on reversed phase C18 OMIX tips (Agilent), all according to the manufacturer’s protocol. Purified peptides were dried, re-dissolved in 20 µl loading solvent A (0.1% trifluoroacetic acid in water/acetonitrile (ACN) (98:2, v/v)) and 5 µL were injected for LC-MS/MS analysis.

### LC-MS/MS and data analysis

LC-MS/MS analysis was performed on an Ultimate 3000 RSLCnano system (ThermoFisher scientific) in line connected to a Q Exactive mass spectrometer or a Q Exactive HF (ThermoFisher scientific). Trapping was performed at 10 μl/min for 4 min in loading solvent A on a 20 mm trapping column (100 μm internal diameter (I.D.), 5 μm beads, C18 Reprosil-HD, Dr. Maisch, Germany) before the peptides were separated on a 150 mm analytical column packed in the needle (75 µm I.D., 1.9 µm beads, C18 Reprosil-HD, Dr. Maisch). Prior to packing of the column, the fused silica capillary had been equipped with a laser pulled electrospray tip using a P-2000 Laser Based Micropipette Peller (Sutter Instruments). Alternatively, peptides were separated on a 500 mm long micro pillar array column (µPAC™, PharmaFluidics) with C18- endcapped functionality. This column consists of 300 µm wide channels that are filled with 5 µm porous-shell pillars at an inter pillar distance of 2.5 µm. With a depth of 20 µm, this column has a cross section equivalent to an 85 µm inner diameter capillary column. Peptides were eluted from the analytical column by a non-linear gradient from 2 to 55% solvent B (0.1% FA in water/acetonitrile (2:8, v/v)) over 30, 60 or 90 minutes at a constant flow rate of 250 or 300 nL/min, followed by a 5 min in 99% solvent B. Then, peptides were eluted by a 5 min in 99% solvent B. The column was then re-equilibrated with 98% solvent A (0.1% FA in water) for 15 min. In the virotrap experiment, the mass spectrometer was operated in positive and data-dependent mode, automatically switching between MS and MS/MS acquisition for the 5, 10 or 12 most abundant ion peaks per MS spectrum. Full-scan MS spectra (400-2,000 m/z) were acquired at a resolution of 70,000 (at 200 m/z) in the orbitrap analyzer after accumulation to a target value of 3E6 for a maximum of 80 ms. The 10 most intense ions above a threshold value of 1.7E4 were isolated in the trap with an isolation window of 2 Da for maximum 60 ms to a target AGC value of 5E4. Precursor ions with an unassigned, or with a charge state equal to 1, 5-8, or >8 were excluded. Peptide match was set on “preferred” and isotopes were excluded. Dynamic exclusion time was set to 50 s. Fragmentation were performed at a normalized collision energy of 25%. MS/MS spectra were acquired at fixed first mass 140 m/z at a resolution of 17,500 (at 200 m/z) in the Orbitrap analyzer. MS/MS spectrum data type was set to centroid. The polydimethylcyclosiloxane background ion at 445.12002 was used for internal calibration (lock mass) in addition to the ion at 361.14660 corresponding to a tri-peptide of asparagine that was spike-in the MS solvents as described in (Staes et al., 2013). Similar settings were used for the GST pull down and immunoprecipitation experiments.

Data analysis was performed with MaxQuant (version 1.6.3.4) using the Andromeda search engine with default search settings including a false discovery rate set at 1% on the PSM, peptide and protein level. All spectral data files were searched with MaxQuant against all human proteins in the Uniprot/Swiss-Prot database (database release version of January 2019 containing 20,413 protein sequences (taxonomy ID 9606), downloaded from www.uniprot.org). For virotrap, the 32 recorded spectral data files were searched together and the search database was complemented with the GAG, VSV-G, and eDHFR protein sequences. For the GST pull down experiments, three different searches were performed, one search for each cell line. For the IP experiments, three different searches were performed, two searches for HEK293T cells treated with interferon-α for 24h and 48h, and one search for HeLa cells treated with interferon-β for 24h. For GST pulldown and IP experiment, each search comprised six spectral data files. The mass tolerance for precursor and fragment ions was set to 4.5 ppm and 20 ppm, respectively, during the main search. Enzyme specificity was set as C-terminal to arginine and lysine (trypsin), also allowing cleavage at arginine/lysine-proline bonds with a maximum of two missed cleavages. Variable modifications were set to oxidation of methionine (sulfoxides) and acetylation of protein N-termini. Matching between runs was enabled with an alignment time window of 20 minutes and a matching time window of 1 minute. Only proteins with at least one peptide were retained to compile a list of identified proteins. In virotrap, 1,214 proteins were identified in all 32 samples. In the GST pulldown, 599 proteins were identified in the experiment with THP-1 cells, 800 with HeLa cells and 933 with HEK293T cells. In the IP experiments, 401 proteins were identified in the experiment with HEK293T cells treated with interferon-α for 24h; 3,795 proteins were identified in the experiment with HEK293T cells treated with interferon-α for 48h; 1,419 proteins were identified in the experiment with HeLa cells treated with interferon-β for 24h. Proteins were quantified by the MaxLFQ algorithm integrated in the MaxQuant software. A minimum of two ratio counts from at least one unique peptide was required for quantification. Further data analysis was performed with the Perseus software (version 1.6.2.3) after loading the proteinGroups table from MaxQuant. Hits identified in the reverse database, only identified by modification site and contaminants were removed and protein LFQ intensities were log2 transformed. Replicate samples were grouped, proteins with less than three or four valid values in at least one group were removed and missing values were imputed from a normal distribution around the detection limit to compile a list of quantified proteins. In the virotrap experiment, 613 proteins were quantified. In the GST pull down experiments, 595 quantified proteins in the experiment with THP-1 cells, 415 with HeLa cells and 933 with Hek293T cells. In the IP experiments, 223 proteins were quantified in the experiment with HEK293T cells treated with interferon-α for 24h; 2,362 proteins were quantified in the experiment with HEK293T cells treated with interferon-α for 48h; 547 proteins were quantified in the experiment with HeLa cells treated with interferon-β for 24h. On the quantified proteins, for each ISG15 bait a t-test was performed for a pairwise comparison with the control condition to reveal specific ISG15 interaction partners. The results of these t-tests are shown in the volcano plot in Fig. 1 and Supplementary Fig. 1-3. For each protein, the log2 (ISG15/control) fold change value is indicated on the X-axis, while the statistical significance (-log p value) is indicated on the Y-axis (Table S1). Proteins outside the curved lines, set by an FDR value of 0.001 and an S0 value of 2 in the Perseus software, represent specific ISG15 interaction partners. The mass spectrometry proteomics data for the virotrap and the GST pulldown experiments have been deposited to the ProteomeXchange Consortium via the PRIDE partner repository with the dataset identifiers PXD018345 and PXD018346.

### Generation of RNF213 knockout mice

B6J-RNF213em1Irc mice were generated using the CRISPR/Cas9 system. Synthetic Alt-R® CRISPR- Cas9 crRNA (Integrated DNA Technologies) with protospacer sequences 5’ CAGAGCTTCGGAACTTTGCT 3’ and 5’ TGTGCCCCTCATCAACCGTC 3’ were duplexed with synthetic Alt-R® CRISPR-Cas9 tracrRNA (Integrated DNA Technologies). cr/tracrRNA duplexes (100 ng/µl) were complexed with Alt-R® S.p. Cas9 Nuclease V3 (500 ng/µl) (Integrated DNA Technologies). The resulting RNP complex was electroporated into C57BL/6J zygotes using a Nepa21 electroporator with electrode CUY501P1-1.5 using following electroporation parameters: poring pulse = 40V; length 3.5 ms; interval 50 ms; No. 4; D. rate 10%; polarity + and transfer pulse = 5V; length 50 ms; interval 50 ms; No. 5; D. rate 40%; polarity +/-. Electroporated embryos were incubated overnight in Embryomax KSOM medium (Merck, Millipore) in a CO2 incubator. The following day, 2-cell embryos were transferred to pseudopregnant B6CBAF1 foster mothers. The resulting pups were screened by PCR over the target region using primers 5’ AGTTTCTTGATCTCTTCCCC 3’ and 5’ CTCCTCCGTCAGATCCCTA 3’. PCR bands were Sanger sequenced to identify the exact nature of the deletion. Mouse line B6J-RNF213em1Irc contains an allele with a deletion of 2854 bp (chr11+: 119440493-119443346) in exon ENSMUSE00000645741 resulting in a frameshift and premature stopcodons (Supplementary Fig. 5).

